# Transcription dosage compensation does not occur in Down syndrome

**DOI:** 10.1101/2023.06.07.543933

**Authors:** Samuel Hunter, Robin D. Dowell, Josephina Hendrix, Justin Freeman, Mary A. Allen

**Affiliations:** Molecular, Cellular, and Developmental Biology, University of Colorado Boulder, 80301 Boulder, USA; BioFrontiers Institute, University of Colorado, 80309 Boulder, USA; Linda Crnic Institute for Down Syndrome, 80045 Aurora, USA; Crnic Boulder Branch, 80309 Boulder, USA; Computational Bioscience, the University of Colorado Anschutz Medical Campus, Aurora, CO, USA, 80309 Boulder, USA; Center for Genes, Environment and Health, National Jewish Health, Denver, CO, USA, 80309 Boulder, USA

**Keywords:** Down syndrome, GRO-seq, RNA-seq, DNA-seq, dosage compensation

## Abstract

**Background:** Trisomy 21, also known as Down syndrome, describes the genetic condition of having an extra copy of chromosome 21. The increase in DNA copy number has led to the "DNA dosage hypothesis", which claims that the level of gene transcription is proportional to the gene’s DNA copy number. Yet many reports have suggested that a proportion of chromosome 21 genes are dosage compensated back towards typical expression levels (1.0x). In contrast, other reports suggest that dosage compensation is not a common mechanism of gene regulation in Trisomy 21, providing support to the DNA dosage hypothesis.

**Results:** In our work, we use both simulated and real data to dissect the elements of differential expression analysis that can lead to the appearance of dosage compensation even when compensation is demonstrably absent. Using lymphoblastoid cell lines derived from a family of an individual with Down syndrome, we demonstrate that dosage compensation is nearly absent at both nascent transcription (GRO-seq) and steady-state RNA (RNA-seq) levels.

**Conclusions:** Transcriptional dosage compensation does not occur in Down syndrome. Simulated data containing no dosage compensation can appear to have dosage compensation when analyzed via standard methods. Moreover, some chromosome 21 genes that appear to be dosage compensated are consistent with allele specific expression.

## Introduction

Trisomy 21 (T21, also known as Down syndrome) is the most prevalent aneuploidy in the human population[1]. All individuals with Down syndrome have a number of standard physical features and some degree of intellectual disability. In addition, they have an increased risk for specific health problems such as congenital heart disease and a decreased risk of others, including solid tumor formation (see review [2]). This altered risk profile arises primarily from the effect of higher levels of transcription of chromosome 21 genes [3, 4, 5]. The increase in DNA copy number and transcription has led to the DNA dosage hypothesis: gene transcription levels are proportional to DNA dosage [6, 7].

Dosage compensation is any mechanism that modulates gene expression to compensate for increased DNA dosage [8]. The most famous and well-studied dosage compensation mechanism involves X inactivation to balance sex chromosome expression levels[9]. In contrast, no dosage compensation mechanism is known to exist for an entire mammalian autosomal chromosome[8]. In Down syndrome, an autosomal chromosome is triplicated, thus, there is tremendous interest in whether dosage compensation exists for any genes on chromosome 21.

Numerous studies of gene expression in Down syndrome cells have reported gene expression levels that do not strictly follow DNA dosage [10, 11, 12, 13, 14, 15]. Others have contradicted these findings, arguing instead that most genes follow the expected 1.5-fold increase, with only a few genes showing lower than expected expression levels[3, 5]. In part, the differences in opinion arise from how each study defines dosage compensation. For example, a permissive definition regards every gene below the DNA dosage expected 1.5 median fold change as dosage compensated[15], but this effectively ignores normal statistical variation inherent to these measurements. A more principled approach looks for deviations from the 1.5 median fold change that exceed statistical expectation[5]. Regardless of the methodology used for identifying dosage compensation, all studies identify at least a small number of genes expressed below expectation, suggesting they may be dosage compensated.

Here we sought to identify the sources of apparent dosage compensation, both technical and molecular. Thus we first focus on short read sequencing data and the statistical methods of assessing differential expression. All differential expression studies use analysis tools, such as DESeq2, to determine the list of differentially expressed genes[16]. These pipelines provide a systematic, reproducible methodology which seeks to maximize the number of accurate differential gene calls while minimizing false positives by accounting for the underlying noise inherent to these data. These tools are exceptionally well designed to discover differential gene expressions when the data fit the expectation of the tool (see review [17]).

However, when used with the default parameters, the underlying assumption is that these differential expression tools work equally well on trisomy data. Yet to date, no study has examined how the presence of an extra copy of chromosome 21 impacts the typical differential expression analysis pipeline. Therefore, we first dissect the typical analysis pipeline to identify issues that could lead to erroneous identification of dosage compensation. To this end, we created simulated transcription data sets for both a disomic (D21) and trisomic individual where no dosage compensation is present by design (i.e. all chromosome 21 genes measure at the expected 1.5x change). Using the simulated data, we modify the typical differential expression analysis pipeline to accurately account for the trisomy nature of the data. We then apply our trisomy-aware analysis pipeline to biological data, both steady state RNA-seq and nascent transcription data, generated from a family where one child has Down syndrome. Our study finds that only a few chromosome 21 genes are expressed or transcribed at lower than the 1.5x expectation.

We hypothesize that the few genes with lower than expected levels may reflect individual alleles with lower-than-expected transcription levels. To address this question, we leverage both the family structure and DNA sequence to determine whether the set of genes with lower-than-expected transcription levels can be explained by allelic variation. Research suggests inter-individual variation is a more substantial contributor to differential expression in T21 studies than sex or aneuploidy status[5, 3]. Consistent with this, tremendous variability in expression levels exists within the population of typical, diploid humans[18]. For example, large-scale studies of gene expression among typical humans find that 83% of genes are differentially expressed between subsets of individuals[19]. Expression quantitative trait loci (eQTL) studies seek to identify loci genome-wide that contribute to observed variation in gene expression levels[20, 21]. We find that most of the genes with lower-than-expected transcription arise because of genetic variations that lead to lower expression levels (known eQTLs).

Our findings show no dosage compensation at either transcription or steady-state RNA levels. RNA-seq analysis pipelines, when not trisomy aware, can lead to the appearance of dosage compensation when there is none. Furthermore, our work shows that natural genetic variation can explain lowly expressed genes that appear to be dosage compensated. Finally, based on our simulated data sets, we create guidelines for accurate differential expression analysis in trisomy cells, which we call trisomy-aware transcription analysis.

## Results

### A naive analysis suggests technical issues in dosage compensation detection

We sought to identify the sources of apparent dosage compensation, both technical and molecular. To this end, we focus on lymphoblastoid cell lines derived from a family of individuals where one child has Down syndrome (Figure 1A). We reasoned that if dosage compensation did occur, it would either occur via inhibition of transcription or via an increase in the RNA degradation of that transcript. Thus we examined both steady-state RNA levels via RNA-seq and nascent transcription with global run-on sequencing (GRO-seq), in triplicate (See Methods and Materials, Supplemental Table 1 for sequencing information, Supplemental Figs. 1 and 2).

**Figure 1.**
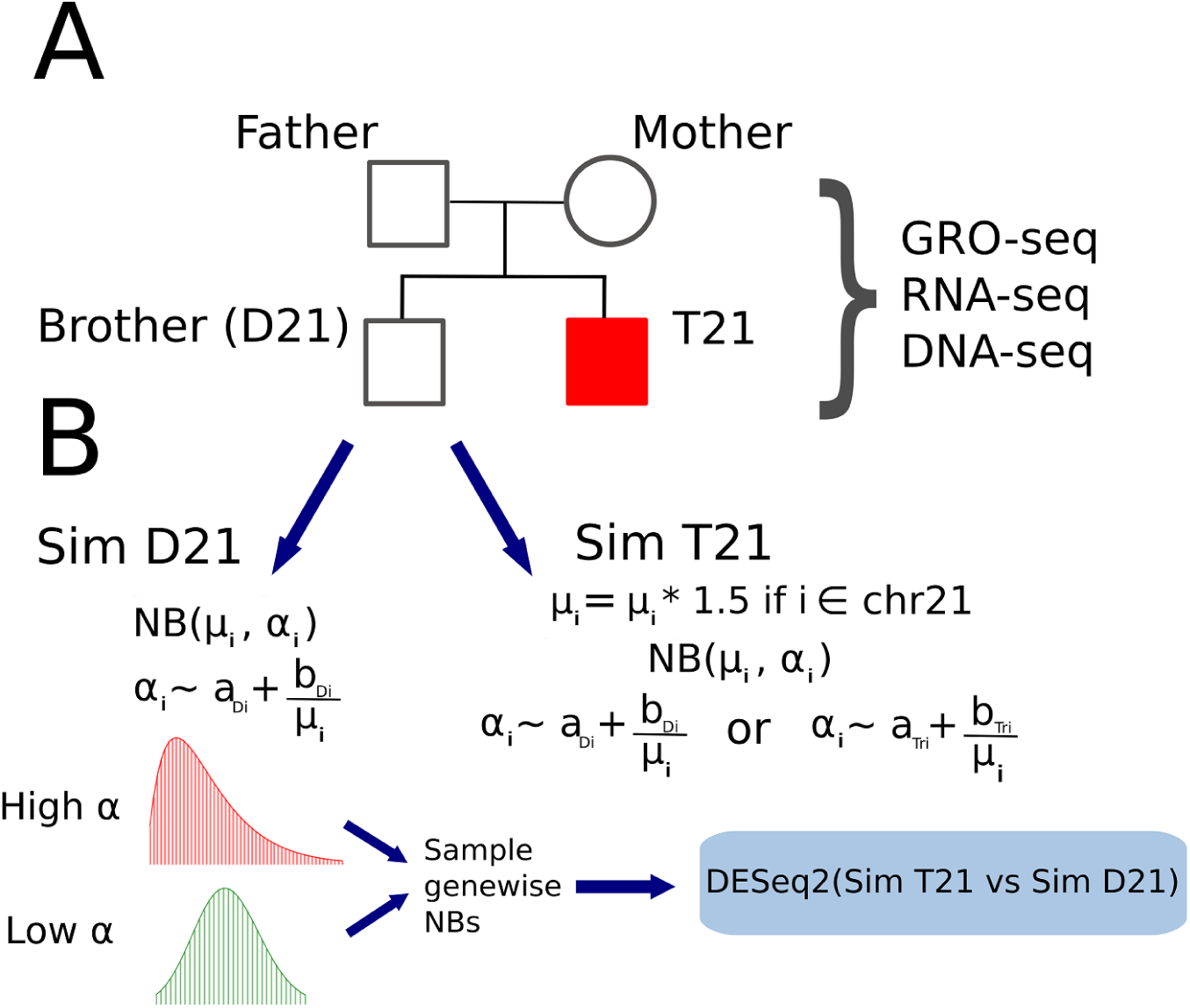
Summary of cell line generation and dataset simulation. (A) Pedigree depicting the relationship of our samples. Lymphoblastoid cell lines (LCLs) were derived from each of the individuals. Libraries for GRO-seq, RNA-seq, and DNA-seq were generated from these cell lines for downstream analysis (B) Simulations generated from the D21 child. The RNA-seq datasets from this individual were averaged together to inform the mean counts (mu) for each gene i. The hyperparameters a (termed asymptotic dispersion) and b (termed extra-poisson noise) are used to inform the genewise dispersion of each negative binomial (NB) distribution. New read datasets for each gene were then generated by random variate sampling from these distributions. For trisomic genes, the mean of the negative binomial distribution (represented as mu) is first multiplied by 1.5, ensuring that calculated fold change estimates between trisomic and disomic genes should yield an expected distribution around 1.5, modulated by dispersion. Varying hyperparameters were used to generate multiple simulated datasets.

As an initial baseline, we first examined the typical differential analysis pipeline that leverages DESeq2[22]. In this naive analysis, we make no adjustments to the defaults inherent to programs within the pipeline. Using the naive approach, we find the median fold change (MFC) of all genes on chromosome 21 in RNA-seq is 1.41, with 57.6% of individual genes having a fold change below the expected 1.5 fold change. The trends in GRO-seq are similar with an overall chromosome 21 median fold change of 1.38 with 48.8% below the expected 1.5 fold change (Supplemental Fig. 3).

To identify genes as dosage compensated, we must specify specific criteria – i.e. how much below the expected 1.5x levels is unusual? First, we considered a fold change cutoff. In this idealized case, if two populations of genes exist (i.e. dosage compensated and not compensated) one would expect to see two distinct but overlapping distributions of fold change in the data. To examine this, we generated a cumulative distribution function of fold change for all chromosome 21 genes in RNA-seq and GRO-seq (Supplemental Fig 3, red line). We noted that the majority of genes were distributed around 1.5; however, no apparent secondary population was immediately clear – instead, we observe a smooth distribution of fold changes below 1.5, suggesting fold change is distributed on a continuum. While this distribution does not in and of itself disprove the existence of transcription dosage compensation, these results argue that there is no obvious cutoff for identifying dosage-compensated genes. Indeed, these results are in agreement with Hwang et. al. findings that dosage compensated genes (defined in their approach as FDR q-value < 0.01) are rare in trisomy RNA-seq datasets[5]. This led to the question of how trisomy data influences differential expression and the assessment of dosage compensation. To address this question, we turn our attention to dissecting the typical analysis pipeline, using DESeq2 as a representative technique, in order to identify how T21 influenced these results.

### Simulations reveal the technical basis of reduced fold change calculations in trisomic datasets

To carefully assess the impact of trisomy data on the standard differential expression analysis pipeline, we need to know a priori the correct answer. To achieve this, we simulated T21 and D21 datasets, using the real data from the child with D21 data as a reference. Briefly, we used the D21 data along with parameters (such as variance) utilized by DESeq2 to create an artificial gene counts table (Fig 1B, see Materials and Methods for full details). The simulated T21 individual was generated in the same way, but now all genes on chromosome 21 are at a 1.5x increase from the simulated D21 individual.

We then run the standard analysis pipeline (Figure 2A) on the simulated data, calculating the fold change and its significance for all genes from each simulation. First, we run the simulation using the parameters obtained from the naive analysis pipeline (Figure 2B) and even though the chromosome 21 genes in the simulated data are at 1.5x, we observe distributions and median fold changes similar to the biological data (MFC = 1.4). Next we seek to isolate individual components of the differential expression pipeline (Figure 2A) by running the simulation several times, modifying the count table’s read depth, replicate number, or variance to test each variables’ effect on differential expression and fold change estimation. For simplicity, we present only the distributions for chromosome 21 (expected to be at 1.5x) and chromosome 22 (expected to be at 1x; all other disomic chromosomes showed the same results as chromosome 22).

**Figure 2.**
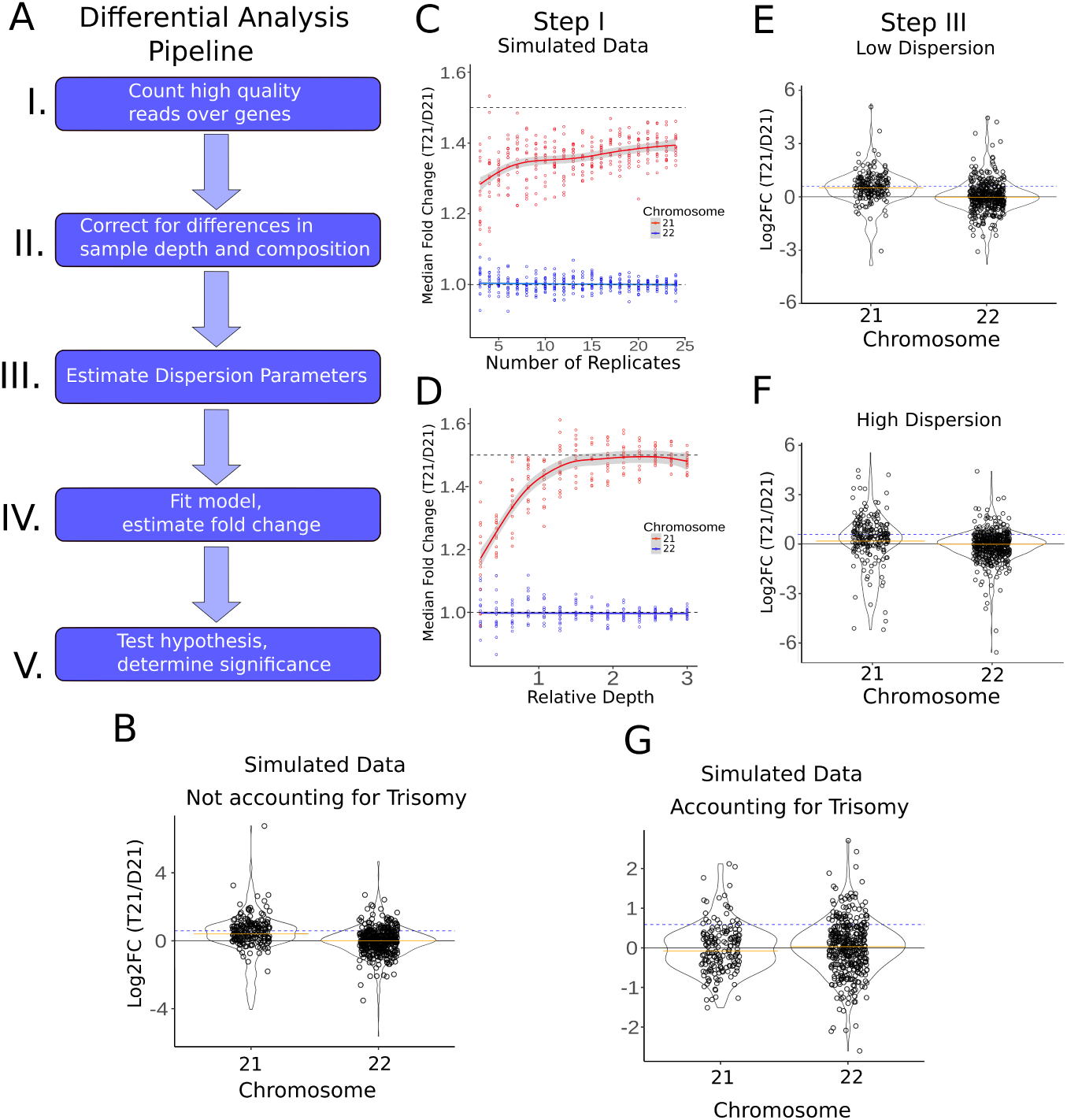
Fold Change distributions of RNA-seq and GRO-seq datasets. (A) Pipeline of differential analysis. Variations at any step have the potential to increase or decrease fold change calculations for chromosome 21 genes (See also Supplemental Fig 10,11,7) (B) Naive differential analysis of simulated T21 and D21 datasets using the similar dispersion parameters found in real data (asymptotic dispersion = 0.03, extra-Poisson noise=3.5, see Supplemental Fig7). For chromosome 21 genes (which were simulated at 1.5x), the median fold change is 1.40. (C) Effects of shifting parameters of simulated datasets. Simulated datasets with varying numbers of replicates (asymptotic dispersion=.01, extra-Poisson noise=1). (D) Simulated datasets with varying levels of depth (asymptotic dispersion=.01, extra-Poisson noise=1). (E) Violin plots showing fold-changes of simulated datasets when dispersion parameters are low (asymptotic dispersion=.01, extra-Poisson noise=1). (F) Violin plots showing fold-changes of simulated datasets when dispersion parameters are high (asymptotic dispersion=.05, extra-Poisson noise=30). (G) Simulated data violin plots showing fold-changes after applying adjustments for each step in the pipeline. Results are consistent with no dosage compensation in T21 datasets in the simulated data.

#### Sequencing depth and read counting methodologies

Tools such as DESeq2 seek to quantify within-condition variability using principled models of read-count data to determine changes between conditions that exceed expected variability[16]. As such, the key first characteristics of sequencing data is the number of replicates available per condition. Therefore we first examined the impact of replication, assuming initially that all replicates are of relatively high quality (low variance). To this end, we simulated data with varying replication, between 2 and 25 replicates per condition. We found that the median fold change estimation generally increased with additional replicates (Fig 2C), consistent with the notion that additional replication is always a good strategy.

The parallel concern to replication is sequencing depth. It is important to note that nascent transcription typically has lower overall counts per gene than RNA-seq when the two protocols are sequenced to roughly equivalent depths because a much larger fraction of the genome is transcribed than is stable. Consistent with this, we noticed that low fold change estimates correlated with low expression levels (Supplemental Fig. 4) in actual data. Therefore we next simulated datasets with varying depths, ranging from 0.1 times to 10 times the depth of the original datasets. We found that decreasing the depth of the simulated datasets resulted in a decreased fold change estimation for many genes, with concomitant reduced median fold change estimates (Fig 2D). Thus it is important to have adequate depth for accurate gene fold change estimates. However, additional sequencing is not an option when reanalyzing public data sets and in some cases increased sequencing depth can be cost-prohibitive. Importantly, increased depth is not expected to fully alleviate the fold change estimation issue; as with increased depth, unexpressed genes are more likely to have reads assigned to them due to noise. Thus we suggest users employ a minimum coverage filter to remove low signal genes as potential false positives when determine which genes are potentially dosage compensated.

In these initial simulations, we noticed that genes with the most dramatic apparent dosage compensation in our naive analysis disappeared in the simulations. Specifically, highly expressed chromosome 21 genes with many genomic repeats, such as ribosomal genes, often appeared at typical expression levels in our real datasets (Supplemental Fig. 5), but not in our simulated data. This suggests that repeat regions shared between chromosome 21 and other chromosomes are sensitive to the mapping strategy. During mapping, these reads can be sponged away from chromosome 21 genes, resulting in a lower fold change estimation. This is, of course, dependent on the employed mapping strategy and how multiple mapped reads are handled. Combining a minimum read cutoff and masking repeat regions or removing multimapping reads effectively removed many of these genes as false positive dosage compensations. We suggest that users filter these genes from their list before dosage compensation analysis, either by masking repeat regions before counting reads or manually removing genes with a high number of genomic repeats before subsequent analysis[23].

#### Size factor calculation for sample normalization

After counting reads, the next step is to normalize the data between libraries (Step II in Fig 2A). Normalization accounts for differences in sequencing depth between samples and is crucial to proper differential analysis. DESeq2 utilizes a median-of-ratios method to find a normalizing “size factor” for each library[22]. In short, DESeq2 assumes that the majority of genes are similarly expressed from sample to sample. By calculating the ratio between the counts for a gene in one sample versus the mean count in all samples and then finding the median of these ratios, samples can be effectively normalized to the genes most likely to remain unchanged in all samples. We thus wondered if this normalization method was being influenced by the trisomic condition of one of our samples, where we expect a priori for a set of genes to be changed between the two samples simply because they reside on the aneuploid chromosome.

To investigate the impact of trisomy on size factor estimation, we removed chromosome 21 from both the actual and simulated data sets. In both cases, chromosome 21 genes had only a minimal effect on size factor calculation (Supplemental Fig 6), consistent with the relatively small proportion of genes on chromosome 21 (approximately 1%) compared to the rest of the genome. Importantly, this result was robust to overall sequencing depth, which we showed by modulating the sequencing depth of our simluations. While the empirical result suggests including chromosome 21 genes in size factor estimation has little effect on the results, we nevertheless recommend removing chromosome 21 genes for the size factor calculation, as this is more consistent with the theory behind the median-of-ratios method.

#### Dispersion estimation and sample replication

We next investigated the effects of the model fitting process of DESeq2 on fold change estimation in T21 cells. In general, gene expression is estimated by fitting a negative binomial distribution with two parameters: the mean and the dispersion (both of which inform the variance of the distribution). Both values can be inferred directly by maximum likelihood estimation, which calculates the mean expression level for each group of replicates, and then determines the dispersion value. However, this method is susceptible to error at lowly expressed genes or when a low number of replicates are available, as systemic noise begins to dominate[22]. More optimal methods, such as DESeq2, employ a Bayesian process allowing information sharing across multiple genes and replicates. In particular, DESeq2 assumes that genes with similar expression levels exhibit similar dispersion values. Information can thus be shared across genes and samples to more accurately estimate gene-wise dispersion, which increases the fidelity of fold change estimation and dispersion calls even within noisier data. However, if this assumption fails – i.e., if a cluster of similarly expressed genes has higher dispersion than expected – the resulting fits will not accurately reflect the underlying biology. We thus endeavored to answer this question: does the presence of a T21 sample affect the model estimation steps of differential analysis? The default method used in DESeq2 for calculating gene-wise dispersion (Step III in Fig 2A) involves starting with the maximum likelihood estimate of dispersion, plotting these values against expression levels, and then fitting an asymptotic curve in the form of *y* = *a* + *b/x* [22]. Here, *a* and *b* are the fitting parameters, *y* is the dispersion estimate, and *x* is the gene expression level. The parameters *a* and *b* control the two ends of the curve; for low-expression genes, the parameter *b* will be more meaningful, leading to higher dispersion estimates for the fitted curve at these genes, a phenomenon we refer to as extra-Poisson noise. For high-expression genes, the *b/x* term trends toward 0, and thus the fitted curve will asymptotically approach the value of parameter *a*, a trend we refer to as asymptotic dispersion. The resulting fitted curve is then used to inform one more round of dispersion estimation, effectively shrinking gene-wise dispersion towards the fitted curve. An increase in either parameter increases the value of the initial dispersion estimate. Still, the effects are felt asymmetrically depending on the expression level and the amount of information available to each gene (i.e., replicates and sequencing depth).

To determine the effects of T21 on dispersion estimation, we extracted the fitting parameters used in dispersion estimation from our biological data sets, with and without chromosome 21 genes. As a control, we also compared removing an equivalent number of random genes from the other chromosomes (see Materials and Methods). We observed an increase in both fitting parameters relating to gene-wise dispersion when trisomic genes were included (Supplemental Fig 7), leading to the question: how does this increase in the initial dispersion fit affect differential calls and fold change estimates?

We sought to quantify and demonstrate how the increase in dispersion would affect differential analysis for genes on chromosome 21; as such, we generated data sets sampled from a negative binomial distribution with the same means but varying dispersion values for chromosome 21 genes in the T21 simulations. When we compared across a wide range of dispersion estimates, we noted higher dispersion parameters resulted in decreased fold change estimations for chromosome 21 genes (Fig 2E, F; Supplemental Fig 8). Specifically, the distribution of fold-change drastically shifted towards 1.0, albeit with a broader spread (MFC = 1.28). The effects of higher dispersion on fold change estimation could be partially offset in simulated data by adding more replicates and depth (Supplemental Fig. 9), consistent with the fact that more replication and read depth are critical to overcoming the effects of systemic noise. Notably, the fact that a high number of replicates creates a MFC closer to 1.5x may also contribute to the disparate results regarding dosage compensation reported in previous studies; large scale studies that integrate several available data sets result in more confident fold change estimation indicating no dosage compensation (i.e. most genes are at the expected 1.5x fold change)[5].

#### Fold change shrinkage and hypothesis testing

The final steps of differential expression analysis are fold change estimation and hypothesis testing. Here, DESeq2 provides the option to utilize a Bayesian method for fold change estimation, using a prior distribution centered around a fold change of 1.0, which effectively shrinks fold change estimates towards 1.0. As with dispersion estimation, the resulting shrinkage effect is more substantial for low-expression genes. These estimates (known as maximum a posterior or MAP estimates) are generally considered more reliable than MLE calculations of fold change for low expression genes [22]. However, this assumption can fail if the prior distribution does not represent the underlying biology, as is true for trisomic genes (Supplemental Fig 10). Researchers who utilize MAP estimates will thus note more genes that appear dosage compensated, although this apparent “compensation” is mainly due to the shrinkage effects of the prior distribution. In general, we recommend users exercise caution when interpreting MAP estimates as evidence for dosage compensation; genes that experience strong fold-change shrinkage should be filtered out from analysis, or MLE calculations should be used instead.

Hypothesis testing in the standard differential analysis pipeline uses the default null hypothesis that each gene’s expression levels are equal in both groups. Given that chromosome 21 genes exist in three copies and DESeq2 seeks to identify deviations from typical, one might expect most of chromosome 21 to be differentially expressed, e.g. statistically significant. However, when we ran our simulation using parameters similar to the real data (Fig 2B, padj < .01), no genes on chromosome 21 are called as statistically significant. This arises because while the median fold change of chromosome 21 encoded genes is elevated (MFC = 1.4), the dispersion genome wide is high enough that this change is not deemed significant. In our simulations, we altered dispersion over a broad range (Supplemental Fig. 8) and while the median fold change varied (Fig 2E,F), no genes on chromosome 21 were deemed statistically significant. In contrast, increasing replicates and read depth, even with higher dispersion values, did yield 126/262 differentially expressed chromosome 21 genes in RNA-seq simulations (MFC = 1.43) (Supplemental Fig 9). In all cases, significant genes showed fold change estimates near or greater than the expected 1.5 fold. Thus, with high replication DESeq2 detects the typical 1.5x transcription levels of chromosome 21 as statistically significant, but finds no genes with lower than expected expression levels.

Arguably, this arises because differential expression analysis is a distinct question from identifying dosage compensation. The default null hypothesis used in hypothesis testing is incorrect for identifying dosage compensated genes, as all of chromosome 21 is expected to be elevated. Consequently, to identify dosage compensation using the standard differential expression pipeline it is necessary to either adjust the null hypothesis or the input data.

The first method is to change the fold-change threshold for the hypothesis tests of chromosome 21 genes from the default value of 1 to the dosage-informed value of 1.5, such that significant genes calls deviate from the DNA dosage-informed expectation. Under these tests, significant gene calls below a fold change of 1.5 are potential candidates for dosage compensation. In our actual and simulated datasets, most chromosome 21 genes are not considered significant when using this method (Supplemental Fig 11). In other words, even for genes with a fold change below 1.5, only significant calls can be interpreted as potentially dosage compensated. However, we note that this method cannot reliably utilize the MAP estimates of fold change, as the prior distribution does not reflect the new alternative hypothesis.

The second method, which we prefer, is to perform an additional normalization step before the differential analysis. In DESeq2, the ploidy number of each gene in each sample can be loaded into a normalizing matrix. The resulting read counts are normalized both by the library’s size factor and the gene’s ploidy number. Subsequent fold change shrinkage and hypothesis testing can thus utilize the default parameters, as even trisomic genes are expected to exhibit a fold change of 1.0 under these conditions (Fig 2G, Supplemental Fig. 12). Thus, significant genes with a fold change of less than 1.0 are candidate dosage-compensated genes. Given we simulated data with a 1.5-fold change, this approach correctly finds no significant chromosome 21 genes in our simulated data, regardless of the dispersion parameters used (Supplemental Fig 13).

In biological data, ploidy normalization brings the distribution of chromosome 21 fold change estimates in line with other chromosomes (Fig 3A-C, Supplemental Fig. 14). Thus chromosome 21 genes exhibited an MFC of 0.96 with dosage normalization in RNA-seq (Fig 3C). Furthermore, 1009 genes were considered differentially expressed between the two brothers, of which only five were on chromosome 21 (out of 143 total chromosomes 21 genes, after read count filtering). In GRO-seq analysis, chromosome 21 genes had an MFC of 0.97 and 3820 genes were differentially expressed, of which 20 genes fell on chromosome 21 (out of 144 total chromosome 21 genes, after filtering) (Supplemental Fig 15). We note that the proportion of significant genes on chromosome 21 is similar to those obtained when we compare two disomic individuals; the distribution of fold-changes and the number of differential genes are consistent so long as the reads are normalized by the ploidy number (Supplemental Fig 16). Normalizing by ploidy is additionally advantageous, as MAP estimates of fold change can be utilized for visualization and downstream analysis. Furthermore, subsequent power analyses are more relevant to trisomic data once the counts have been adjusted[24].

**Figure 3.**
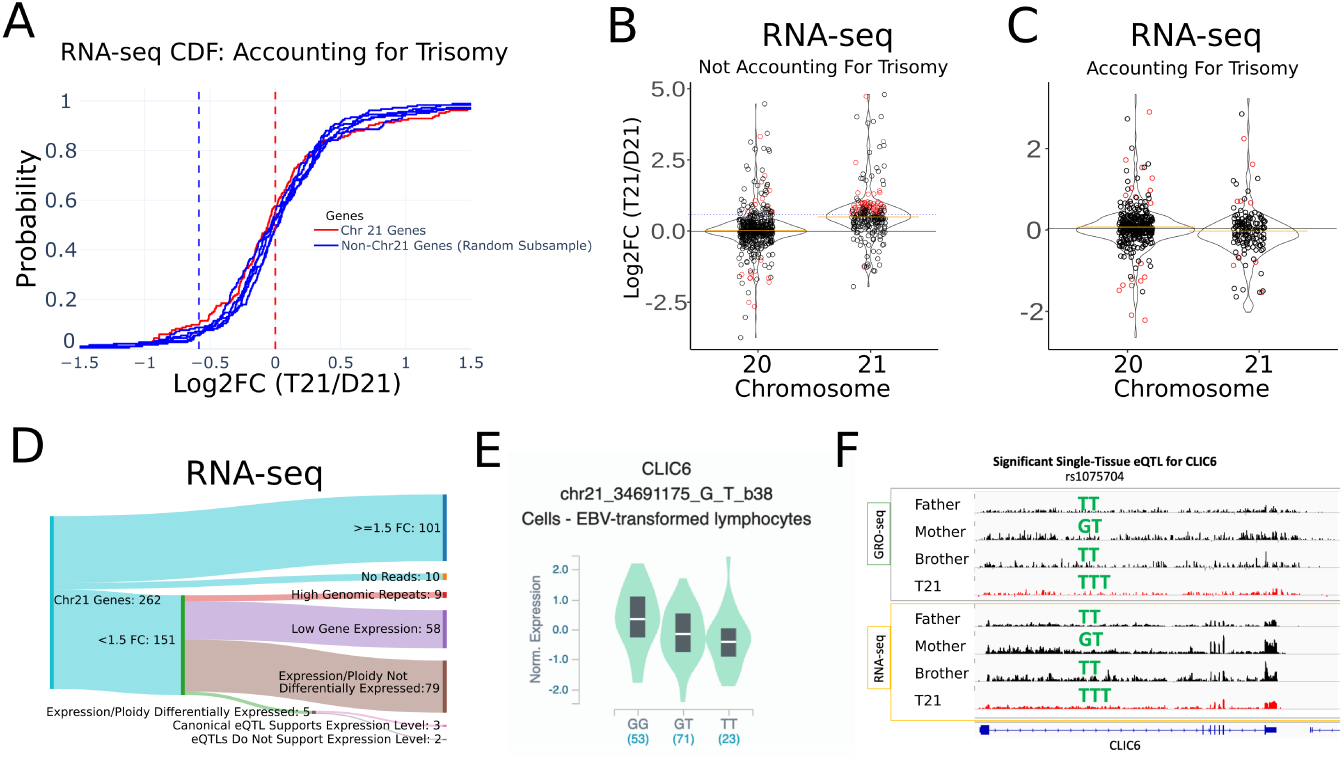
Alternative explanations to disparate fold change estimates. (A) Cumulative distribution plot of fold changes found in real RNA-seq data, after accounting for trisomy. Solid red line indicates all chromosome 21 genes. Each solid blue line is a randomly selected set of genes from all other chromosomes. (B) Violin plots indicating Log2 Fold Change between T21 and D21 samples. Significant gene calls are colored red (padj < .01). (C) Same as (B), but using a trisomy-aware pipeline for analysis. (D) Sankey diagram depicted the filtering process of our RNA-seq analysis. The initial 151/262 genes identified as potentially dosage compensated can alternatively be explained by genomic repeats, high variance from low expression genes, or technical artifacts related to failing to normalize the data to the ploidy number. Remaining genes can be explained by the presence of eQTLs (See also Supplemental Figs 17)). (E) Example boxplot indicating relative expression of the gene CLIC6 with one eQTL. (F) Genome viewer tracks for the gene CLIC6 for all four family members, in GRO-seq (top) and RNA-seq (bottom). The T21 track is indicated in red. The allelic makeup of the eQTL in (E) is indicated by the green text above each track.

**Figure 4.**
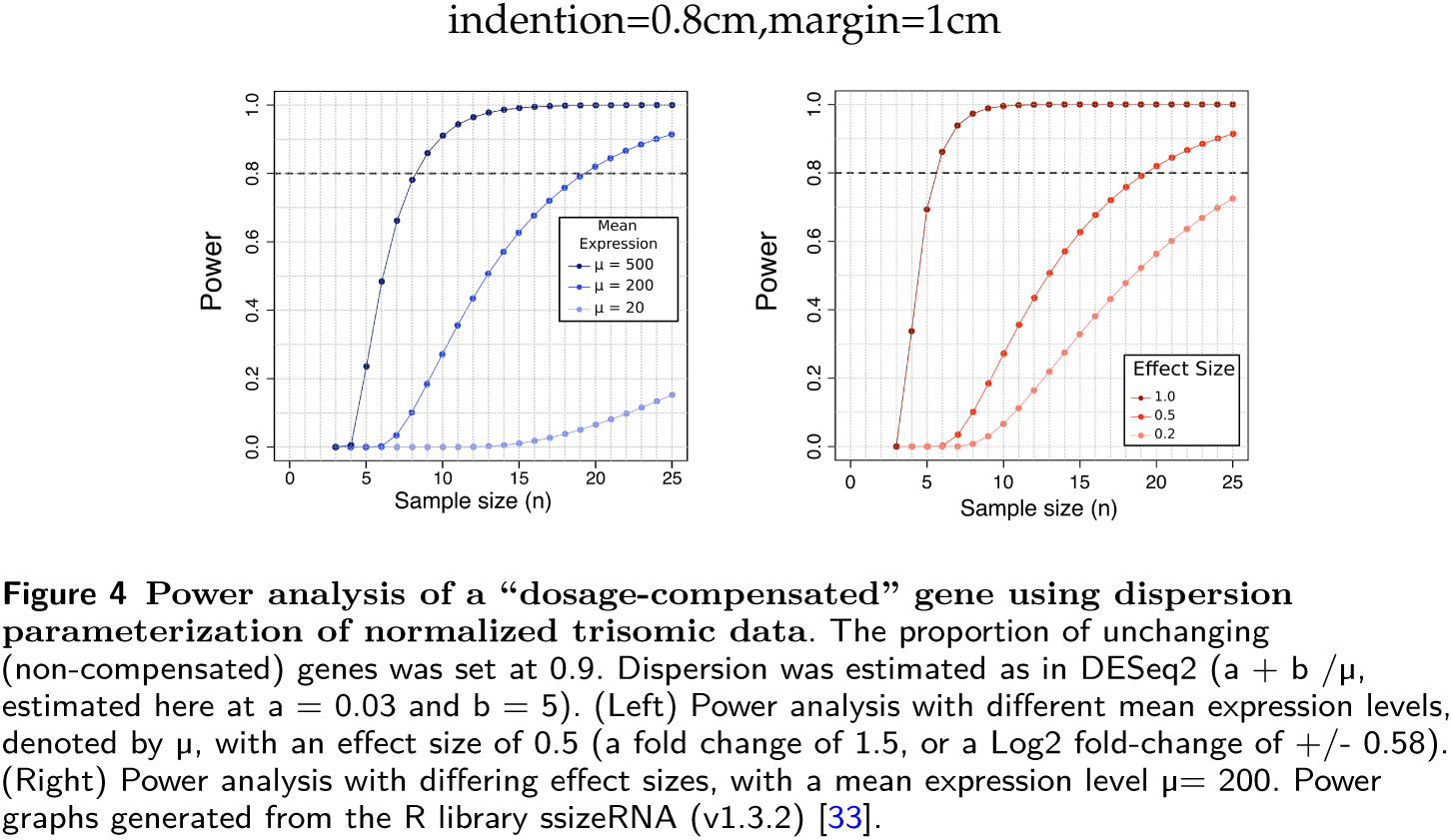
Power analysis of a “dosage-compensated” gene using dispersion parameterization of normalized trisomic data. The proportion of unchanging (non-compensated) genes was set at 0.9. Dispersion was estimated as in DESeq2 (a + b /μ, estimated here at a = 0.03 and b = 5). (Left) Power analysis with different mean expression levels, denoted by μ, with an effect size of 0.5 (a fold change of 1.5, or a Log2 fold-change of +/- 0.58). (Right) Power analysis with differing effect sizes, with a mean expression level μ= 200. Power graphs generated from the R library ssizeRNA (v1.3.2) [33].

**Figure 5.**
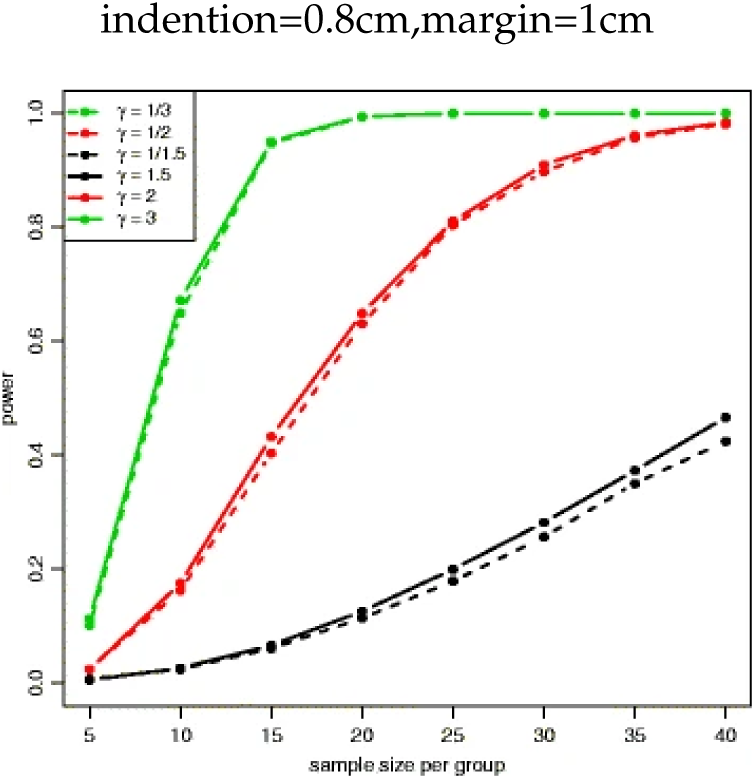
Reproduced from Yu et al.: Effect size versus power in differential expression analysis. Power plot at *μ*=100 for the Wald test with equal dispersion parameters. Power was calculated at 8 different sample sizes and 6 different fold changes under the alternative hypothesis with *μ*=100 and *α*=0.001. Power is higher for larger sample sizes and higher absolute fold changes (Figure caption text and image from [24]).

**Figure 6.**
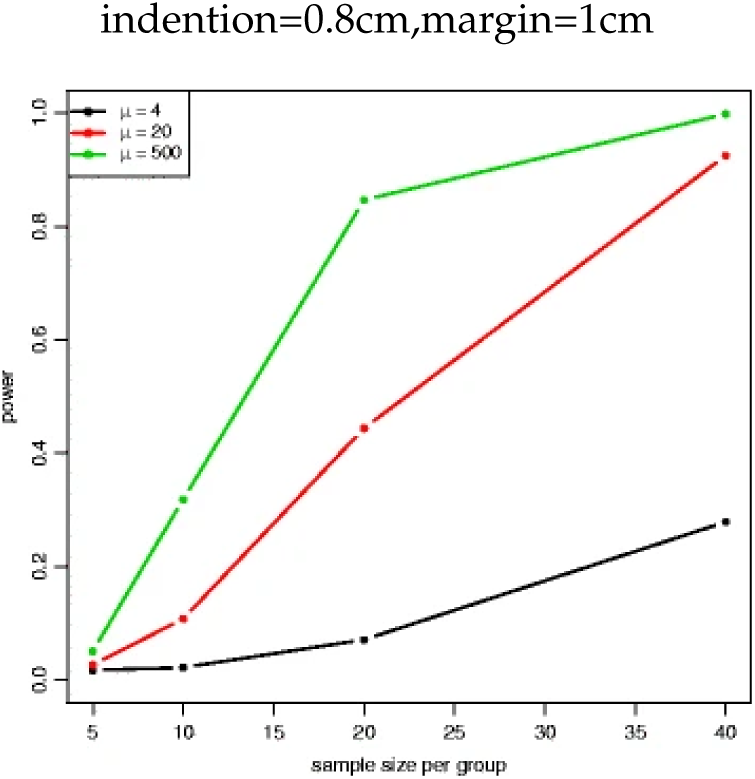
Reproduced from Yu et al.: Effect of means and dispersion on power calculation. Power plot at *γ*=2 for the Wald test with unequal dispersion parameters. Power was calculated at 4 different sample sizes and 3 different expression levels with *γ*=2 under the alternative hypothesis and *α*=0.001. Power is higher for larger sample sizes and higher expression levels(Figure caption text and image from [24]).

In all, our simulations led us to the construction of a differential analysis pipeline more suited to trisomic samples. First, we find that multi-mapped reads can cause a handful genes to appear dosage-compensated; we thus suggest removing these reads prior to analysis. Next, we note that low expression genes are especially prone to appearing dosage compensated; we thus suggest employing a minimum coverage filter to avoid these false positives. Finally, our trisomic samples appeared to be noisier relative to their disomic counterparts; this variance resulted in lower fold change estimations for many chromosome 21 genes. As such, we suggest either employing a normalization matrix to adjust all gene counts by their ploidy number, or using the appropriate null hypothesis to determine potential dosage compensation in trisomy. Rather than relying on cutoffs, our trisomy-aware pipeline uses DESeq2’s significance calls to determine which gene expression levels are significantly different than their expected values based on gene dosage (Fig 3C,D, Supplemental Fig 17). Furthermore, by accounting for the genes which contribute the most variance in fold-change calculations, we remove likely false positives from the analysis. We thus contend that our pipeline is more rigorous determinant for dosage compensated genes which properly accounts for the underlying variance of the data.

### Reduced fold change on chromosome 21 are consistent with identified eQTLs

After adjusting the data analysis pipeline to be trisomy-aware, we conclude that nearly all chromosome 21 encoded gene transcription levels (GRO-seq) and expression levels (RNA-seq) are proportional to DNA dosage(Supplemental Fig 13). However, a small number of genes on chromosome 21 remain potentially dosage compensated, as precisely five genes in RNA-seq and 20 genes in nascent transcription (Fig 3D, Supplemental Fig 15) have lower-than-expected levels.

Given the small number of genes remaining and the lack of known dosage compensation mechanisms in humans, we reasoned that there might be a genetic basis for the reduced expression of these genes. More concretely, we hypothesize that apparent dosage compensation could arise from allele-specific sequence variation. To explore this possibility, we sought to compare the genome sequence of these individuals to known expression quantitative trait loci (eQTL) where a specific sequence variant is associated with lower expression within the population[25]. Thus we performed whole genome DNA sequencing on the cell lines of all four family members (see Materials and methods for full details; Supplemental Table 1 for sequencing information).

We used the GATK package to call SNPs in each member of the quartet[21]. Briefly, reads were mapped and realigned before variants were called using Haplotype caller (see Methods and materials for full description). Because chromosome 21 is triploid in the individual with Down syndrome, Haplotype caller was used twice— once with the default ploidy of two and once with ploidy set to three. Variants called using the ploidy of three version were kept only for chromosome 21 in the individual with Down syndrome, otherwise the default calls were used.

We reasoned that any genetic variation that reduces chromosome 21 gene expression in the general population could be present in our T21 sample, leading to reduced expression levels in the individual. To test this hypothesis, we compared genome variations identified in each individual to previously identified eQTLs in the GTEx database[21, 26]. We limited our search to eQTLs identified in either lymphoblastoid cell lines or their nearest related tissue (Whole Blood). For this analysis, we required the variant to be identifiable in at least one of the parent samples as well as the child with Trisomy 21. In RNA-seq, we found that the reduced expression of three of the five apparently dosage-compensated genes (CLIC6, ITSN1, C2CD2) could potentially be explained by known expression controlling polymorphisms (Fig. 3D, Supplemental Table 2). For example, the rs1075704 eQTL exists in the human population as a GG, GT, or TT (Fig 3E), and the T allele correlates with lower expression of CLIC6 in lymphoblastoid cells. This SNP, rs1075704, shows variable CLIC6 gene expression across the disomic samples in both our RNA-seq and GRO-seq (Fig. 3F). In our trisomic sample, CLIC6 has an expression level less than 1.5x the average of all disomics, with the genotype TTT. So the lower than expected expression observed at CLIC6 can be explained by the genotype, which is not considered by the typical differential expression pipeline. Consistent with the allele identity, these three genes also have lower than expected transcription in GRO-seq.

We also reasoned that the eQTL data in GTEx may be incomplete, that there may exist other alleles that lead to reductions in expression data. To explore this possibility, we next compared the two parents to each other, reasoning that differences observed between the parents could be subsequently inherited by either of the two children. Using this technique, we found that 9 of the remaining 17 genes in GRO-seq were also differentially transcribed between the parental samples, suggesting the lower than expected levels in the trisomy sample may be modulated by an inherited parental haplotype (Supplemental Fig 17). Altogether, we found reasonable explanations for most genes (60%) which fell below expected levels in T21 (Fig 3). These results were consistent in both RNA- and GRO-seq (Supplemental Fig 17).

## Discussion

We sought to add clarity to the conflicting reports in the literature concerning whether molecular dosage compensation occurs within individuals with Down syndrome[5, 10, 12, 13, 14, 11, 15]. Our results uniformly suggest that dosage compensation in T21 is rare, if not completely absent, in transcriptomics data, both nascent and steady-state RNA. Using simulated data, we found that computational pipelines developed for disomic samples could lead to erroneous conclusions on trisomy data, and this likely contributes to confusion in the literature. We were able to use our simulated data to create a trisomy-adjusted differential expression analysis pipeline that correctly estimates the simulated fold change of chromosome 21 genes. When we applied this modified pipeline to our actual samples, most of the apparent dosage compensation was lost. Importantly, the remaining genes with lower than expected expression were predominantly attributable to alleles with reduced expression levels. Thus our work agrees with other recent studies suggesting no reduction in RNA expression levels via dosage compensation in T21[5] and provides explanations regarding previous reports to the contrary. In addition, we have extended the conclusion of no dosage compensation to nascent RNA transcription.

Many analysis pipelines (including the popular DESeq2 algorithm) use a null hypothesis that the fold change between two samples is 1, and are thus the default analysis is ill-equipped for fold change estimation and differential analysis when DNA dosage suggests an alternative fold change is expected. These native settings can be easily adjusted, leading to a more reliable analysis. In summary:

1. Apply a minimum read coverage filter, depending on read depth (30 used in this study)
2. Mask repeat regions or remove multi-mapping reads for read counting
3. Remove chromosome 21 genes for size factor calculation
4. For noisy samples, increase sequencing depth or replication
5. Adjust the null hypothesis or normalize read count by ploidy number

While we focused here on dosage compensation, our findings have implications for using trisomic samples in differential expression analysis. We have shown that the inclusion of trisomic samples in a differential expression analysis pipeline can inflate the dispersion of the data if not adequately accounted for. The presence of trisomic samples in a data collection can affect parameter estimation within the differential analysis pipeline, even when the trisomic sample isn’t part of the final comparison. For example, comparing the son with D21 to his father results in two sets of significant gene calls, depending on whether the trisomic sample is included in the upstream processes (Supplemental Fig 16).

We also note that, by design our study did not compare large groups of individuals with and without T21. In any study, genes below expected expression levels could arise from trisomy 21-altered pathways, molecular dosage compensation or genetic allele frequency differences. We used a family of related individuals to minimize allele variation and focus specifically on molecular dosage compensation. We found no molecular dosage compensation but we did find allele variation that could be driving lower-than-expected expression levels. In a larger cohort of unrelated individuals with Down syndrome, some alleles may have lower-than-expected expression levels. Consistent with this, a recent study of large groups of unrelated disomic and trisomic individuals found many genes are expressed below a 1.5-fold change expectation [27].

As there are many genes with sequence variations that lead to higher or lower expression[26, 21], our work leads to speculation about whether highly expressed alleles could be selected against in a Down syndrome background. More than 75% of trisomy 21 embryos are lost due to spontaneous abortion and it is currently unclear why[27]. If genes on chromosome 21 exist that are deleterious pre-birth when expressed at high levels, individuals with T21 harboring three copies of these alleles would be more likely to be lost. This would result in allele bias within the population of live individuals with T21, favoring the lower expressed allele. Assessing whether particular alleles have a skewed frequency in the T21 population relative to typical individuals requires large numbers of genomes from both typical and individuals with Down syndrome. As an alternative, an extensive collection of T21 RNA-seq could be leveraged toward identifying allele bias. In recent years, the allelic fold change method has been implemented for quantifying eQTL effects[28]. While this model is currently constrained to disomic samples only (i.e., it only allows for three different allelic combinations), the model could be extended (allow for a fourth allelic state) and applied to trisomic samples. As such, population eQTL data could be used to identify lower-than-expected fold change at some alleles in Down syndrome. It remains to be seen whether any chromosome 21 allele bias exists in individuals with T21 [12].

Finally, we note our work was limited to only the exploration of dosage compensation at the transcription level and does not account for potential changes at the protein level due to aneuploidy. Indeed, recent studies identify several protein complexes which are potentially dosage compensated in Down syndrome, likely due to a change in stoichiometric ratios of their respective subunits[29]. Furthermore, new research contends that a majority of proteins undergo dosage compensation in other aneuploidies, such as those found in common cancers[30]. In any case, mRNA abundance is not fully predictive of protein abundance, and conclusions regarding transcription dosage compensation cannot necessarily be extended to the protein level, or absolute quantification methods used to assess protein abundance.

## Materials and Methods

### Cell Culture of LCLs

The lymphoblastoid cell lines of 4 individuals were obtained from the Translational Nexus Biobank (COMIRB 08-1276), University of Colorado School of Medicine, JFK Partners. Lymphoblastoid cell lines (LCLs) were seeded in upright T-25 suspension flasks with 10 ml RPMI (10% FBS, 1X L-Glutamine, 1X Penicillin/Streptamycin). These were passaged approximately every 2 to 3 days by pelleting the cells via centrifugation (300 x g, 5 min) and resuspension. Cells were grown to an approximate density of 1 million cells per ml, before being harvested for subsequent experiments. The three RNA-seq replicates per sample were based on three separate growths of cells and the three GRO-seq replicates per sample were based on three separate growths. The correlation between replicates can be seen in Supplemental Figure 2.

### Nuclei Isolation

Cell cultures were collected in 50 ml Falcon tubes and centrifuged in a fixed-angle rotor centrifuge at 300 x g, 4*^◦^*C, for 5 minutes. The supernatant was poured off, and 10 ml ice-cold PBS added to resuspend the cell pellet. The previous spin and PBS wash was repeated 2 more times. Cells were then resuspended in 10 ml ice-cold Lysis Buffer (10 mM Tris-HCl pH 7.5, 2 mM MgCl_2_, 3 mM CaCl_2_, 0.5% IGEPAL, 10% Glycerol, 2 U/mL SUPERase-IN, brought to 10 ml with 0.1% DEPC DI-water). The resuspended cells were incubated for 10 minutes on ice. Nuclei were then centrifuged in a fixed-angle rotor centrifuge at 1000 x g for 10 minutes at 4*^◦^*C. The resulting supernatant was poured off, and pellet was resuspended with 1 ml lysis buffer, using a wide-mouth P1000 pipette tip. The volume of was brought to 10 ml with lysis buffer, and centrifuged for 1000 x g, 4*^◦^*C, 5 minutes, in a fixed-angle rotor centrifuge. The lysis buffer wash was repeated as above. Nuclei were then resuspended in 1 ml freezing buffer (50 mM Tris-HCl pH 8.3, 5 mM MgCl_2_, 40% Glycerol, 0.1 mM EDTA pH 8.0, brought to volume with 0.1% DEPC DI-water), and transferred to a 1.7 ml Eppendorf tube. Nuclei were pelleted at 1000 x g, 4*^◦^*C, 5 minutes. Resulting supernatant was removed by pipetting, and pellet resuspended with 500 *µ*l freezing buffer. The nuclei were again pelleted at 1000 x g, 4*^◦^*C, 5 minutes. The supernatant was removed, and the nuclei resuspended in 110 *µ*l freezing buffer, and stored at -80*^◦^*C until library preparation.

### GRO-seq and Library Preparation Methods

Wash solutions for Anti-BrdU-beads were prepared ahead of time; Binding buffer (0.25X SSPE, 37.5 mM NaCl, 0.05% Tween, 1mM EDTA pH 8, 0.2% SuperRNAseIN), Low Salt Wash Buffer (0.25X SSPE, 0.05% Tween, 1mM EDTA pH 8, 0.2% SuperRNAseIN), High Salt Wash Buffer(0.25X SSPE, 137.5mM NaCl, 0.05% Tween, 1 mM EDTA pH 8, 0.2% SuperRNAseIN), TET Buffer(10 mM Tris-Cl pH 8.0, 0.05% Tween, 1 mM EDTA pH 8.0, 0.2% SuperRNAseIN), Elution Buffer (150 mM NaCl, 50 mM Tris-Cl pH 7.5, 20 mM DTT, 0.1% SDS, 1 mM EDTA, 0.2% SuperRNAseIN).

Anti-BrdU beads (Santa Cruz, sc-32323-ac) were prepared by washing and blocking. Per sample, 60µl were washed twice in 500 *µ*l binding buffer. beads were blocked in 500 *µ*l blocking buffer(1X binding buffer, 0.1% PVP, 1 *µ*g/mL BSA UltraPure, 0.002 more superRNAseIN). Beads were then resuspended in 450 *µ*l binding buffer.

Run-on reactions were performed as in [31]. In brief, ice-cold isolated nuclei (100 *µ*L) were added to 37*^◦^*C 100 *µ*L reaction buffer (Final Concentration: 5 mM Tris-Cl pH 8.0, 2.5 mM MgCl_2_, 0.5 mM DTT, 150 mM KCl, 10 units of SUPERase In, 0.5% sarkosyl, 500 *µ*M rATP, rGTP, and Bromo-UTP, 2 *µ*M rCTP). The reaction was allowed to proceed for 10 min at 37*^◦^*C, followed by the addition of 500 *µ*L of Trizol LS. RNA was extracted once with phenol-chloroform, washed once with chloroform, and precipitated with 3 volumes of ice-cold ethanol and 1-2 *µ*L GlycoBlue. The pellet was washed in 75% ethanol before resuspending in 18 *µ*L of DEPC-treated water.

Libraries were prepared similar to [32]. In brief, RNA was treated with 2 *µ*l NEB Fragmentation Buffer at 94 degrees for 5 min. The RNA was then buffer exchanged via BioRad P-30 (or a G-25) column per the manufacturer’s protocol. Next, 2 *µ*l DNaseI and 5 *µ*l of 10X RQ1 DNase buffer and water were added to create a 1X final concentration of the buffer. After incubation at 37*^◦^*C for 10min, 5 *µ*l DNAse stop solution was added, and the reaction was placed at 65*^◦^*C for 5 min. As prepared above, the beads in 450 *µ*l binding buffer was immediately added to this mixture.

Fragmented nascent RNA was purified using Anti-BrdU beads via washing with 500 *µ*l for 1 min with each solution (binding buffer, low salt buffer, high salt buffer, TET buffer). Between washes, beads were spun down per manufacturer instructions. RNA was eluted from the bead via soaking the beads in 125 *µ*l elution buffer at 42*^◦^*C, 10 min 2 times. The beads were then added to acid phenol-chloroform, as was the elute. The RNA was washed with chloroform and precipitated with 5M NaCl and 3x the ethanol column. The second round of Anti-BrdU bead binding and extraction enriched BrdU-labeled products was completed as above. NEBNext Ultra II RNA was used to transform the nascent RNA into an RNA-seq library. The product was amplified 15 *±* three cycles and products >150 bp (insert > 70 bp) were size selected with 1X AMPure XP beads (Beckman) before being sequenced.

### RNA-seq

RNA was isolated from the cells via Trizol extraction. NEBNext rRNA Depletion kit was used to remove rRNA. NEBNext Ultra II RNA was used to transform the RNA into an RNA-seq library. The product was amplified 15 *±* three cycles and products >150 bp (insert > 70 bp) were size selected with 1X AMPure XP beads (Beckman) before being sequenced.

### Mapping and Visualization of RNA datasets

The fastq files for GRO-seq and RNA-seq were trimmed and mapped to the GRCh38/hg38 reference genome and prepared for analysis and visualization through our in-house pipelines. In short, resulting fastq read files were first trimmed using bbduk (version 38.05) to remove adapter sequences, as well as short or low quality reads. Reads were mapped with HISAT2 (version 2.1.0), and resulting SAM files converted to BAM files with Samtools (version 1.8). Multimapped reads were filtered from these files. BedGraph files were generated using Bedtools (version 2.25.0), and converted to TDF files for visualization in IGV using IGVtools (version 2.3.75). Quality metrics were generated with FastQC (version 0.11.8), Preseq (version 2.0.3), RSeQC (version 3.0.0). Figures were generated through MultiQC (version 1.6).

### Differential Expression Analysis

Differential transcription was performed using the DESeq2 (version 1.26.0) R package (R version 3.6.3). Gene counts were generated using featureCounts (version 1.6.2) from the R Subread package (version 1.6.0), counting over the gene body region (+150 from transcription start site to annotated transcription end site) to avoid the 5*^′^* peak. In both RNA-seq and GRO-seq analyses, reads were counted over the gene (including intronic regions), to ensure that results between these two protocols should be comparable. Annotations were downloaded from RefSeq (release number 109, downloaded August 14, 2019 from UCSC genome browser). Only annotations with both RNA-seq and GRO-seq signals were considered, again to keep both analyses comparable. For featureCounts, BED6 region files were converted to SAF format with the following command: awk -F "\t" -v OFS="\t" ‘print{$4, $1, $2, $3, $6}’ region.bed > region.saf. Only the highest transcribed isoform of each gene was considered.

For our corrected analysis, we made use of DESeq2’s normMatrix parameter. The normalization matrix was generated by assigning each gene in the analysis its ploidy number divided by 2. For all genes not on chromosome 21, this number was thus 1. For genes on chromosome 21 samples in trisomy, this number was 1.5. We also removed reads within regions of genomic repeats, set the betaPrior parameter to false, and set an expression level cutoff at the second quintile of the baseMean counts for all genes. River plots were generated using the web interface of the program Sankeymatic (https://github.com/nowthis/sankeymatic.git).

The script and full table of results are available at the github repository listed below.

### Simulation of trisomy and disomy datasets

Reads were simulated using the negative binomial model as part of the scipy (version 1.8.0) Python package (version 3.6.3). The negative binomial means for each gene were estimated by averaging the counts across replicates for the D21 child samples in RNA-seq. For the disomic simulations, these means were used to directly inform the negative binomial parameters. For the trisomic simulations, the means for chromosome 21 genes were first multiplied by 1.5, to simulate the expected increase in dosage. The negative binomial instance is parameterized as follows:

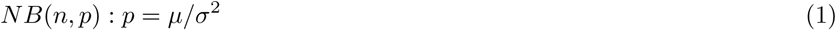

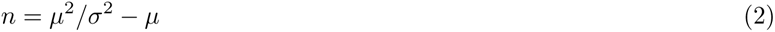

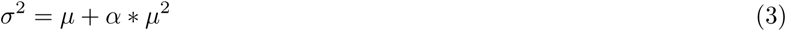

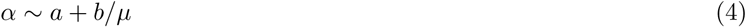

Where *a* and *b* are controllable hyperparameters. For the disomic genes, we used values of .01 and 1 for *a* and *b*, respectively. For trisomic genes, we varied these values from .001 to 1.2 (for *a*) and 1 to 100 (for *b*). The negative binomials were also scaled based on the depth of the original biological samples, from 0.1 times the depth to 3 times the depth. Each simulated counts file was generated with a minimum of three replicates for both the disomy 21 and trisomy 21 samples. For details, see the github repository for these scripts.

### Whole genome sequencing

DNA was prepared using a variety of methods then sequenced on a Highseq 2000 to an approximate depth of 40x (see table fastq for full details). The father, mother, and individual with Down syndrome were first sequenced at Illumina and received as one bam file per person, then split by read group (RG) into individual lane bam files using samtools (version 0.1.19) view. Those files were then converted to fastq using bedtools (version 2.16.2) bamtofastq. All other sample files were received as fastq files. Each fastq file was mapped to hg38 using bowtie(version 2.0.2) with the setting –very-sensitive and using options to retain both library preparation and sequencing reaction information. Tracking sample level information is essential for detecting and removing individual sequencing reaction error rates. SAM files were converted to sorted BAM files using samtools(version 1.2) view and sort. All files for one individual were merged using Picard tools(version 1.72) MergeSamFiles. BAM files were sorted and duplicates were marked using Picard tools(version 1.72) SortSam and MarkDuplicates.

### Whole genome variant calling

GATK (version 3.3-0) was used for variant calling. BAM files were realigned using In-delRealigner with optional flags –known Mills_and_1000G_gold_standard.indels.hg38.vcf -known 1000G_phase1.indels.hg38.vcf [Note: the realignment table was created using all the merged files with RealignerTargetCreator optional flags -known Mills_and_1000G_gold_standard.indels.hg38.vcf -known 1000G_phase1.indels.hg38.vcf]. Then, BaseRecalibrator and PrintReads was used to recalibrate the bases (optional arguments –knownSites Mills_and_1000G_gold_standard.indels.hg38.vcf -knownSites 1000G_phase1.indels.hg38.vcf -knownSites dbsnp_138.hg38.vcf). Haplotypes where called two times, via HaplotypeCaller; once with the optional flags -nct 4 –emitRefConfidence GVCF –dbsnp dbsnp_138.hg38.vcf –variant_index_type LINEAR –variant_index_parameter 128000 and another time with those flags and the flag -ploidy 3. Then vcf files were created for each trio (mother, father, child) using GenotypeGVCFs. To create a vcf that contained chr21 as triploid in only the individual with Down syndrome we created our own code in Python to combine the two types of family vcf files. This program combined the family vcf files by taking any lines that started with “chr21_” or “chr21” from the vcf files with the triploid child variants, and all lines that did not start with “chr21_” or “chr21” from the family diploid vcf files. The vcf file is provided at the Zenodo link listed below.

### GTEx datasets/SNP analysis

We downloaded the GTEx (version 8) database of eQTLs and their associated genes for all available tissues[26]. We merged these databases and filtered out all SNPs identified in our DNA-seq experiments that were not present in the merged GTEx database. We compared the expected effects of each SNP from GTEx with the allelic ratios of each of our datasets. We then asked whether there was at least one SNP that could explain the expression/transcription levels we observed in the real data when each parent was compared to the child with T21. The scripts and full results table are available at the github repository listed below.

### Data and analysis code availability

DNA sequence is available at the SRA with accession number PRJNA966937, RNA-seq and GRO-seq data sets are available within GEO with accession number GSE230999. Genome output and input tables, including vcf are available at doi: 10.5281/zenodo.7527453. RNA-seq dataset mapping information is available at doi: 10.5281/zenodo.7527541. GRO-seq dataset mapping information is available at doi: 10.5281/zenodo.7527537. Analysis output tables, including output of differential expression analysis (naive and corrected) and GTEx comparison table are available in the github along with all scripts and analysis pipelines utilized. The github repository can be found at: https://github.com/Dowell-Lab/DS_Normalization

## Competing interests

Authors MA and RD have a patent for measuring transcription factor activity from eRNA activity. RD was a co-founder of Arpeggio Biosciences. All other authors have no competing interests.

### Author’s contributions

MAA and RDD were responsible for dataset generation, initial concept and analysis, manuscript proofreading, and initial normalization technique. SH was responsible for dataset simulations, final dataset analyses, figure generation, SNP analysis, and drafting the manuscript. JH and JF coordinated genome sequencing efforts.

## Acknowledgements

We would like to thank the employees of the Translational Nexus Biobank (COMIRB 08-1276), the University of Colorado School of Medicine, JFK Partners (Angela Rachubinski, Gessi Pino, Karl Pfenninger, Holly Sullivan). We thank the University of Colorado BioFrontiers Institute Next-Gen Sequencing Core Facility (Amber Sorenson, Jamie Kershner) and Illumina (Linda Gutierrez), which performed sequencing. We acknowledge the BioFrontiers Computing Center at the University of Colorado Boulder for providing High Performance Computing resources supported by BioFrontiers IT (Matt Hynes-Grace, Dan Timmons, Jonathan Demasi). We thank several students/volunteers/research associates that participated in organizing and confirming the results of the genome sequencing: Steve Cape, David Olson, and Philip Cash. Research in was supported by a Sie Post-doctoral Fellowship, NIH R01HL156475, NIH R03HD103995, NIH R01AI156739 and a Linda Crnic Institute for Down Syndrome Seed grant.

## Additional Files

Additional file 1 — Supplemental Figures

Additional file 2 — Supplemental Table 1

Additional file 2 — Supplemental Table 2

article amsmath amssymb,amsthm,enumitem graphicx listings color setspace [english]babel [utf8]inputenc fancyhdr [margin=1in,footskip=0.25in]geometry hyperref changepage bbm [figurename=Response Fig.]caption

xr @colorado.edu1rFig1117Summary of cell line generation and dataset simulation (A) Pedigree depicting the relationship of our samples. Lymphoblastoid cell lines (LCLs) were derived from each of the individuals. Libraries for GRO-seq, RNA-seq, and DNA-seq were generated from these cell lines for downstream analysis (B) Simulations generated from the D21 child. The RNA-seq datasets from this individual were averaged together to inform the mean counts (mu) for each gene i. The hyperparameters a (termed asymptotic dispersion) and b (termed extra-poisson noise) are used to inform the genewise dispersion of each negative binomial (NB) distribution. New read datasets for each gene were then generated by random variate sampling from these distributions. For trisomic genes, the mean of the negative binomial distribution (represented as mu) is first multiplied by 1.5, ensuring that calculated fold change estimates between trisomic and disomic genes should yield an expected distribution around 1.5, modulated by dispersion. Varying hyperparameters were used to generate multiple simulated datasetsfigure.1rFig2218**Fold Change distributions of RNA-seq and GRO-seq datasets**(A) Pipeline of differential analysis. Variations at any step have the potential to increase or decrease fold change calculations for chromosome 21 genes (See also Supplemental Fig 10,11,7) (B) Naive differential analysis of simulated T21 and D21 datasets using the similar dispersion parameters found in real data (asymptotic dispersion = 0.03, extra-Poisson noise=3.5, see Supplemental Fig7). For chromosome 21 genes (which were simulated at 1.5x), the median fold change is 1.40. (C) Effects of shifting parameters of simulated datasets. Simulated datasets with varying numbers of replicates (asymptotic dispersion=.01, extra-Poisson noise=1). (D) Simulated datasets with varying levels of depth (asymptotic dispersion=.01, extra-Poisson noise=1). (E) Violin plots showing fold-changes of simulated datasets when dispersion parameters are low (asymptotic dispersion=.01, extra-Poisson noise=1). (F) Violin plots showing fold-changes of simulated datasets when dispersion parameters are high (asymptotic dispersion=.05, extra-Poisson noise=30). (G) Simulated data violin plots showing fold-changes after applying adjustments for each step in the pipeline. Results are consistent with no dosage compensation in T21 datasets in the simulated datafigure.2rFig3319**Alternative explanations to disparate fold change estimates** (A) Cumulative distribution plot of fold changes found in real RNA-seq data, after accounting for trisomy. Solid red line indicates all chromosome 21 genes. Each solid blue line is a randomly selected set of genes from all other chromosomes. (B) Violin plots indicating Log2 Fold Change between T21 and D21 samples. Significant gene calls are colored red (padj < .01). (C) Same as (B), but using a trisomy-aware pipeline for analysis. (D) Sankey diagram depicted the filtering process of our RNA-seq analysis. The initial 151/262 genes identified as potentially dosage compensated can alternatively be explained by genomic repeats, high variance from low expression genes, or technical artifacts related to failing to normalize the data to the ploidy number. Remaining genes can be explained by the presence of eQTLs (See also Supplemental Figs 17)). (E) Example boxplot indicating relative expression of the gene CLIC6 with one eQTL. (F) Genome viewer tracks for the gene CLIC6 for all four family members, in GRO-seq (top) and RNA-seq (bottom). The T21 track is indicated in red. The allelic makeup of the eQTL in (E) is indicated by the green text above each trackfigure.3 rsup:qc*_p_lots*12**QC metrics of RNA-seq and GRO-seq datasets**

The submitted manuscript by Hunter et al. provides a thorough description of statistical artifacts that arise when analyzing RNA-seq data from aneuploid samples with methods designed for euploid samples. The authors demonstrate that naive use of such tools “off the shelf" has resulted in contradictory and misleading conclusions about autosomal "dosage compensation" in the literature. Using simulation of trisomic gene expression data, they trace the sources of error to specific steps of standard analysis pipelines and present practical guidelines for correcting these biases. Reanalysis of empirical data (both nacent RNA-seq [GRO-seq] and steady state RNA-seq) from trisomy 21 samples demonstated little evidence of dosage compensation, with the small number of genes expressed below the 1.5x expectation better explained by cis-eQTLs on the corresponding haplotypes. This is an important contribution to the field, and the following comments and suggestions are relatively minor:

1) In the Results section, page 3, lines 44-50, I am not sure that hypothesis of dosage compensation requires the existence of two discrete, but overlapping distributions. One could imagine a model where dosage compensation exists on a continuum (though I think the results from this study also disprove that model).

We agree with this point and note that the section in question is meant to tackle previous attempts at defining "dosage-compensated genes". Indeed, the most principled approach is to statistically test the fold change of each individual genes relative to a disomic dataset (which involves no specific fold-change cutoff). As the reviewer points out, the possibility that dosage compensation occurs on a spectrum is mentioned within our paper, but still no dosage compensated genes are identified.

We recognize that the structure of the section in question may have led to this confusion, and have rephrased some of the offending sentences (Results section entitled, "A naive analysis suggests technical issues in dosage compensation detection", third paragraph).

2) What was the basis for selecting the low expression threshold used in your study? Is this threshold still necessary given the later step of adjusting the null hypothesis or including a ploidy normalization factor? I would have thought that low expression would then manifest simply as low power to detect dosage compensation (but would not tend to generate false signal of dosage compensation).

We selected a threshold based on the quintiles’ distributions, per Supplemental Figure 4 and Supplemental Figure 12. Specifically, we chose the 40th percentile for the cutoff, as the distributions past this point were less variant. This cutoff is indeed arbitrary but mirrors similar approaches used for independent filtering in differential analysis[22].

Furthermore, the reviewer is correct; the other adjustments can alleviate this step, per Supplemental Figure 12, but the highly variant genes remain for downstream analysis. A summary statistic (like the median fold change of the chromosome, utilized by this and other studies) is influenced by these lower expression genes even after corrections, so it is still worth removing them for these types of summary applications.

As the reviewer correctly gathered, for the purpose of determining potential dosage compensation via statistical significance, these genes do not show up as significant in either case. Specifically, such lowly expressed genes are generally filtered out via the independent filtering step of DESeq2 or other similar pipelines.

3) In one of the lists of suggestions, the exclusion of multi-mapped reads was mentioned. Was the impact of this actually tested? I would imagine that simply restricting to uniquely mapped reads would present its own biases, as some of the multi-mapped reads in fact originated from the locus in question.

We thank the review for this comment, and the impact of multi-mapping reads is now more clearly discussed in Supplemental Figure 5. The major impact of multimapping reads is on the fold change of the rRNA genes present on chr21. The rRNA genes are highly transcribed from several locations within the genome and therefore, the fold change calculated when including multi-mapped reads is likely incorrect. We have added text to clarify this point.

4) On page 5, line 50-62, it wasn’t obvious to me the direction in which the failure to exclude chromosome 21 during expression normalization would bias results. I realize that the actual impact was small because chromosome 21 contains few genes, but is the direction of the effect that failing to normalize in this way generates erroneous signal of dosage compensation?

We appreciate the reviewers observations on this point and have added additional clarifying information to the text.

In summary, read count normalization excluding chromosome 21 results in a slightly lower size factor for the T21 sample and slightly higher size factors for D21 samples. Including chromosome 21 genes thus slightly shifts the read count distribution of T21 samples lower, and D21 samples higher. While this result indeed would trend towards aberrant dosage compensation, the effect is very small, and both estimates fall well within the confidence interval of the other. We show-case the magnitude and direction of this effect in Supplemental Figure 6.

5) On page 9, line 10, consider a term other than "alternative hypothesis", as I think you are referring to the mis-specification of the null hypothesis.

We thank the reviewer for catching this potentially confusing statement. The reviewer is correct. We have updated the language to reflect the proper terminology.

6) In the discussion, it may be worth discussing literature showing evidence of dosage compensation at the level of translation / protein levels. Are these studies affected by similar biases, or do you believe that dosage compensation is occurring at those downstream stages?

We thank the reviewer for this comment. Many of the proteomic studies employ an absolute quantification approach, which we speculate aren’t subject to the specific sampling/modeling biases we observe here. That said, estimates of RNA-protein correlation levels may be affected, as some of these papers do rely on modeling estimates of fold change from RNA-seq.

We have included a paragraph in the discussion highlighting this point in more detail. The discussion of other studies is also reviewed in the answer to the reviewer’s question 7.

7) The conclusions regarding natural selection in trisomic individuals felt misplaced and was also confusing from an evolutionary genetics standpoint. I would consider removing that paragraph and providing a forward-looking statement that is more closely tied to your findings.

We thank the reviewer for pointing out this confusion in our text. The use of language that is related to evolutionary genetics (population, natural selection) was unintentional. We now avoid those words. The discussion has been altered to explain some of the differences between our study and the other studies in the literature may be due to a bias in the allele frequency in viable T21 individuals.

8) The authors focus much of their effort to highlight the erroneous detection of dosage compensated genes using a non-trisomy aware pipeline for differential expression (e.g. "false-positives"). While I agree that this is a worthwhile highlighting of erroneous conclusions drawn from prior studies, it would also be of major benefit of this study to understand the impact of experimental design (e.g. mean depth, number of replicates) on the power to detect true dosage compensation ("true-positive rate" or "power"). Even if conducted from an in-silico perspective, this would serve to highlight two key points: (1) it is likely that sample sizes of studies attempting to discover dosage compensation were too small in the past to reliably detect these effects and (2) how much more would sample-sizes (e.g. siblings) have to increase to detect a gene with 80% power for dosage compensation? A supplementary plot of sample-size on the x-axis and power to detect true dosage compensation on the y-axis would be a useful addition for scientists attempting to use transcriptomic signatures beyond the trisomy 21 context here as well.

We thank the reviewer for noticing this point. Within the framework of differential expression analysis, interpretations from power analysis are best suited for individual experiments. Practically, differences in basal expression levels of genes, differences in the user-estimated effect size of “dosage compensation”, and differences in dispersion parameterization all influence power from one experiment to another (see Yu et. al. 2017 [24] for a complete discussion). As such, any graph generated using our data would be specific only to our data and may not generalize to future experiments. If this is still of interest to the reviewer, we can include such a graph with the above disclaimers, as shown below:

With that said, there are many excellent studies which explore different avenues for estimating power using the output of DESeq2, and our recommendations for adjusting the differential expression pipeline to account for trisomy are fully compatible with these power analysis pipelines. In the text of the document (Results section, “Simulations reveal the technical basis of reduced fold change calculations in trisomic datasets" subsection “Fold change shrinkage and hypothesis testing" towards the end), we have recommended the study from Yu et al. which explores the many dimensions of power analysis in differential expression studies. For reference, included below are two graphs from their work, simulations that explore the relationship the reviewer requested above (with dispersion parameterization based on Yu et. al. data).

9) The authors have also not commented on the widely used statistics of allelic fold change (aFC) when measuring differential expression. While it is not the immediate subject of investigating using DESeq2 (based on relative read-count abundance), providing a comment in the discussion section on the utility of looking for deviations from expected allelic ratios (e.g. 1:1 in heterozygotes for disomy, 1:2 or 2:1 in disomic heterozygotes) is a way that could provide a robust set of tests for dosage compensation within RNA-seq data. The authors take care to highlight the impact of eQTLs on lowering expression (thereby appearing as dosage compensation), but for genes with a heterozygote that is not affecting absolute expression these ratios of allelic abundance should be a testable hypothesis regarding dosage compensation between disomic and trisomic chromosomes. This does not warrant necessarily performing additional analyses focused on allele-specific expression but should warrant some additional discussion as a potential for alternative statistical evidence for or against dosage compensation in the transcriptome beyond normalized read-counts as in DESeq2.

We appreciate this suggestion from the reviewer, and in response have included in the discussion text on how allelic fold change can be used as an avenue for assessing expression level estimation in trisomy across populations. Importantly, a more detailed attempt at pursuing this approach is best suited to follow-up studies that employ a populations based approach for trisomy gene expression analysis.

## Typos, etc

1. Page 3, Line 59: "Question" is repeated twice.
2. Page 10, Line 6: spelling of "Lymphoblastoid"
3. Page 14: It is not stated whether eQTLs only from a specific tissue were preferred from GTeX or if the eQTL being significant in any tissue was sufficient for this analysis. If some description of the filtering process for tissues could be made here that would substantially help clarify whether the eQTL effects are relevant within the LCL-based context of this experiment. As an alternative, the authors may also consider using the eQTLgen dataset which is a larger dataset based on only whole-blood to avoid potential tissue-specific eQTL from the GTeX project (e.g. Brain, etc). We thank the reviewer for catching these typos. We have clarified which tissues/cells were used (which indeed were only Whole Blood or LCLs specifically) for eQTL comparisons and included a table (Supplemental Table 2) specifically identifying the SNPs, their related genes, the tissues the SNP was identified in, the sample genotypes, and the estimated normalized effect size.
4. Page 15: Please update the URL values of "xxxxxx" for GEO and SRA We apologize for the oversight, the data is available at SRA with the accession number PRJNA966937 and GEO with accession number GSE230999, which is now reflected in the paper.
5. Page 18: If Figure 2 C,D could have the same y-axis scaling that would make the comparisons between these two experimental conditions easier to parse. Similarly for panels Figure 2 E,F if the y-axis could be truncated to [-5,5] that would make these plots more immediately comparable. We have standardized the scaling for the plots above. To avoid removing data on 2F, we instead extended the y-axis of Figure 2E to [-6,6]
6. Figure S12: Should be a reference to Figure 2G?
7. Figure S16: Should be a reference to Figure 3D? We have corrected what was an unfortunate cross referencing error, and these should now be correct.
8. I would appreciate from a reproducibility standpoint if the README in the github repository could be more expository for the scripts that performed each specific analysis (e.g. "script X generated the data represented in Figure 2A"). This would substantially help the replication of the in silico experiments.

We thank the reviewer for this suggestion. We have updated the github with more explanatory information in a README file about which scripts generated which figures.

## Reviewer: 2

Trisomy 21 causes Down syndrome, the most common genetic disease in humans. Despite years of research in Down syndrome, there is substantial debate whether the extra copy of chromosome 21 causes a proportional 1.5-fold increase in transcript levels of the genes located on chromosome 21 or whether dosage compensation attenuates chromosome 21 gene expression. In this manuscript, Hunter et al. present a rigorous and quantitative analysis of the gene expression profiles of cells with trisomy 21. In their study, Hunter et al. conclude that there is no dosage compensation in trisomy 21.

Why do other studies and analyses of trisomy 21 cells conclude that dosage compensation takes place? In this manuscript, Hunter et al. take an elegant approach and simulate data for trisomy 21 with no dosage compensation and analyze it using commonly used tools by the research community. These tools are widely used by bioinformaticians, not considering the biology of the sample. That is, trisomy 21 cells harbor 47 chromosomes compared to 46 chromosomes in the control sample, and the expectation for genes on chromosome 21 is different from the rest of the genome. In the end, Hunter et al. provide a series of recommendations to properly analyze data of cells harboring an extra copy of a human autosome. Lastly, because the authors analyze the gene expression of cells isolated within a family, they can show that allelic-specific expression introduces variability in a handful of chromosome 21 genes whose expression does not seem to increase as predicted.

Overall, the manuscript is well written. The analysis and conclusions reached by Hunter et al. will impact the Down syndrome research community and the general scientific community as gene expression analysis is broadly used to do research.

Minor comments:

Page 3 line 59, the word question is duplicated.

We thank the review for catching this typo, it has been corrected.

## Reviewer: 3

As the authors correctly point out, potential gene dosage compensation in Down syndrome (DS), and in other aneuploid situations, is a topic with general importance. Here, the authors have carried out (i) simulations of RNA-seq datasets from D21 and T21 samples, and (ii) actual RNA-seq, and GRO-seq, from lymphoblastoid cell lines established from a family including a DS proband. There analyses, particularly for the simulated datasets (including varying numbers of technical/biological replicates), are comprehensive and potentially valuable. However, based on my reading of the manuscript (including the Methods section, figures, and supplementary data) it appears that the actual (non-simulation) part of the study lacks technical replicates. Probably for this reason, conventional and useful visualizations of the RNA-seq results, such as by volcano plots and heatmaps, are not included in the current version of their manuscript.

The authors should validate the findings of their simulated datasets by carrying out 3 technical replicates of their RNA-seq (at least the RNA-seq, if not the GRO-seq). Technical replicates appropriate (and standard) for this type of work would be independent RNA preparations from separate wells or separate flasks of each LCL, followed by preparation and sequencing of 3 libraries for each LCL (each individual in the family). Since one library has already been made and sequenced, two more should now be made and sequenced. Then, the authors will be able to show "zoomed-out" overviews of their primary data, especially volcano plots, with associated statistical criteria.

All this being said, if I have MISSED something in the paper and in fact the technical replicates HAVE been done, then I’d be happy to look at it again, if the authors can more clearly describe their technical replicates, and provide the requested data overview figures.

We apologize for not being more clear on the number and type of replicates we gathered. Data was generated in biological triplicates, now described more clearly in the methods section (“Cell Culture of LCLs”). In general, we avoided MA plots or volcano plots as these “zoomed-out" plots are of limited use when discussing fold changes over a single chromosome. However, we have updated the manuscript with a comparison of the correlation of the replicates in Supplemental figure 2.

## Reviewer: 4

The study of Hunter et al is useful and should be published, particularly because of the GRO-seq data. The conclusions are justified by the experimental evidence and analyses of the data. However, the MS is too long for its message and could be reduced to about half of its present length.

We concur with the reviewer that the manuscript could be shorted for an analytical audience. However, we sought to not only identify the sources of false dosage compensation but also to do so in a manner that was generally accessible – even to the less initiated.

In addition, metaphorical or "literary" language should be avoided. Some examples: "raises red flags", "off the shelf", "healthy samples"…

We thank the reviewer for this comment and have removed the metaphorical phrasing to be more exact and descriptive.

The "teaching" sentence in the abstract could be eliminated "we recommend all researcher working …".

We have removed the sentence in question.

The Discussion does not contain comparisons of the content of this MS with those in the literature, but rather a repeat of the current work of the authors.

We thank the reviewer for this observation and have added more context to the discussion. Specifically, we have added discussions of different approaches for fold change estimation in trisomy, including a request from a different reviewer to discuss allelic fold change and translational dosage compensation.

