## Supplementary material for "Transcription dosage compensation does not occur in Down syndrome": Supplmental

### Differential Expression Analysis in Trisomy 21

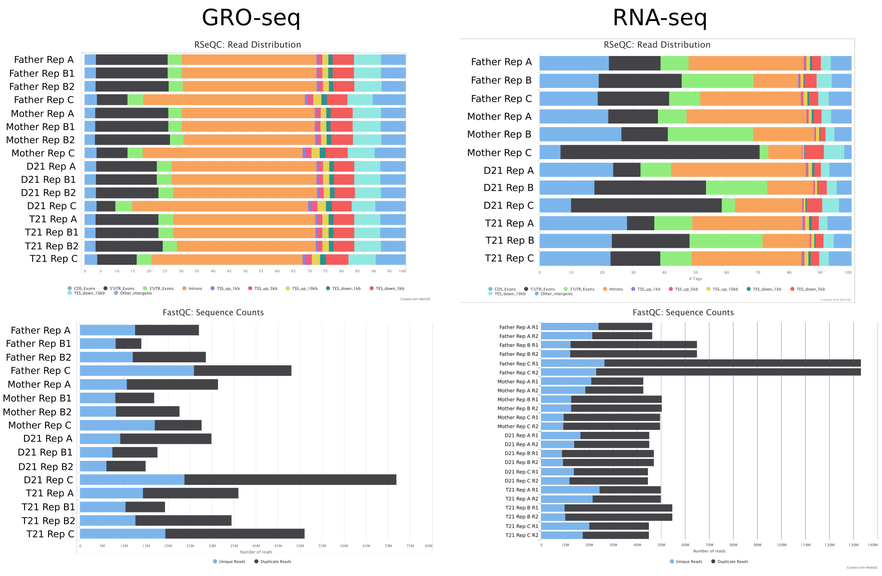


Supplementary Figure 1: **QC metrics of RNA-seq and GRO-seq datasets.** (Top) RSeQC read distribution plots of all datasets, showing relative abundance of reads at each listed genomic feature. (Bottom) FastQC plots showing number of total reads, and proportion of duplicate reads for each dataset.

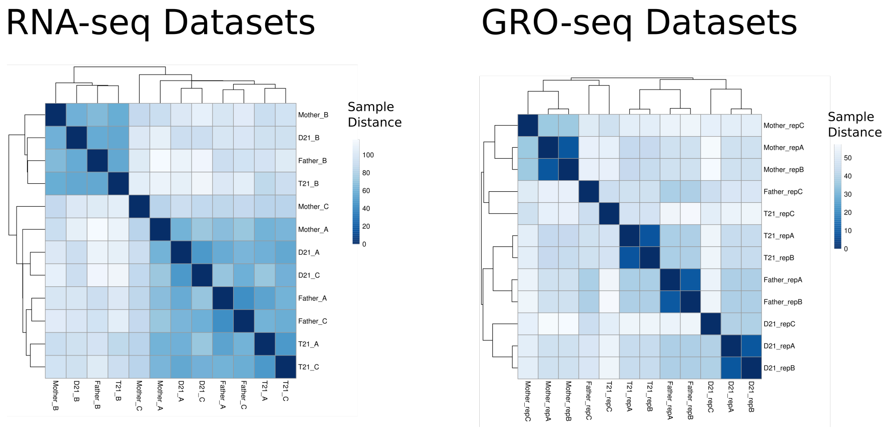


Supplementary Figure 2: **Sample Distance between GRO-seq and RNA-seq datasets** Euclidean distance between replicates (Left) RNA-seq datasets (Right) GRO-seq datasets. The letters A, B, C refer to biological replicates. For more information on samples, see the methods section.

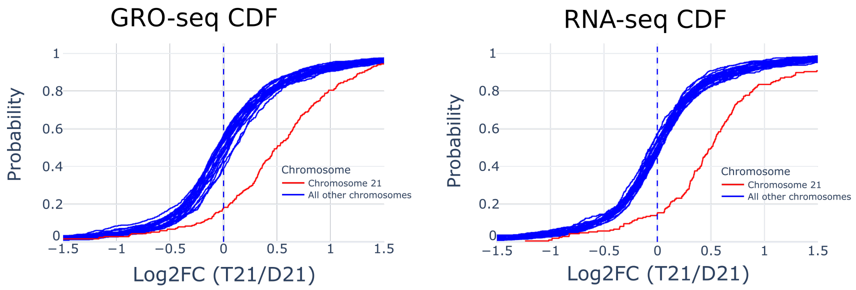


Supplementary Figure 3: **Cumulative distribution plot of fold changes of each chromosome.** All diploid chromosomes in blue; chromosome 21 in red. Dosage compensated genes are often identified using a fold change cutoff, indicated by each vertical dotted line (red:1.5x fold-change, blue: 1,0x fold-change); however, the reduced fold change estimations of these genes can also be explained by biological or technical variance. Left: GRO-seq. Right: RNA-seq.

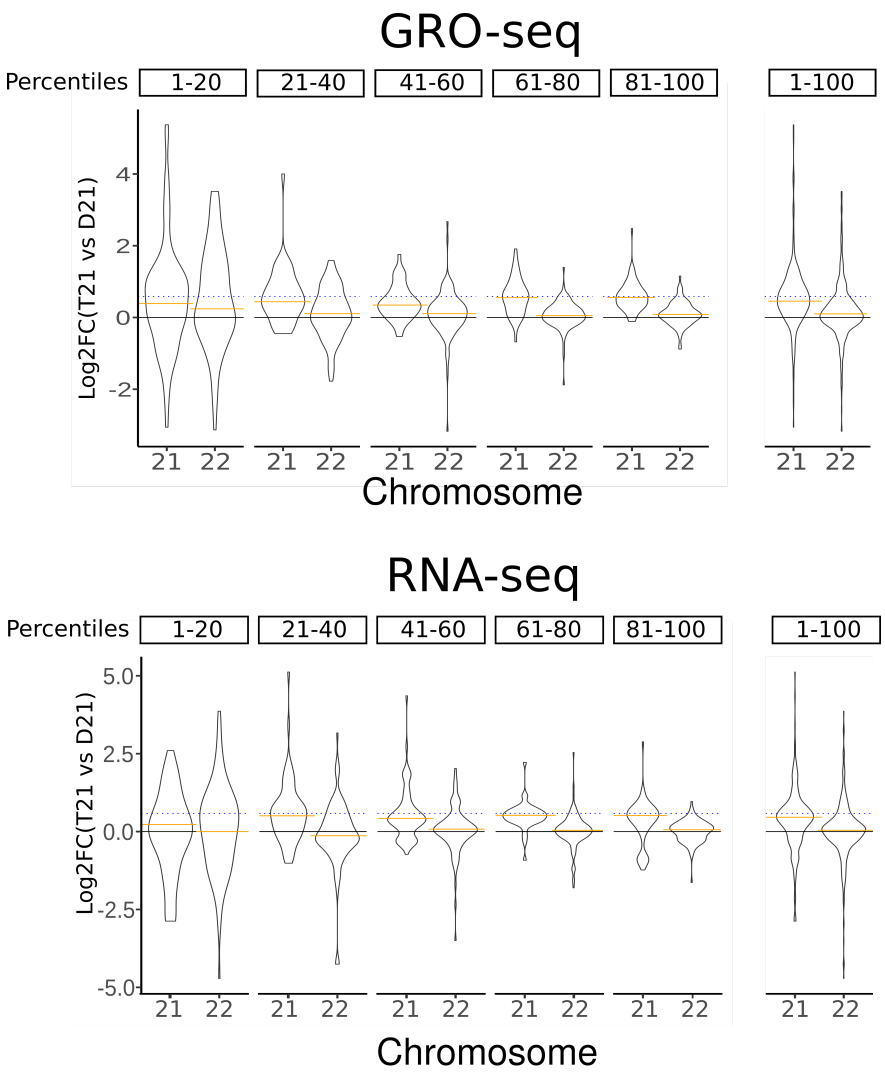


Supplementary Figure 4: **Fold change estimations (T21 vs D21) across expression levels in GRO-seq (top) and RNA-seq (bottom).** Genes are grouped by expression quantiles. Lower quantiles, on the left, contain the genes with the lowest expression (and therefore the genes with the highest technical variability in measurements). The lowest expression quantile has a noticeably lower fold change estimate for chromosome 21 genes. On the far right (labeled 1-100), the figure shows all genes in one violin plot regardless of expresion levels.

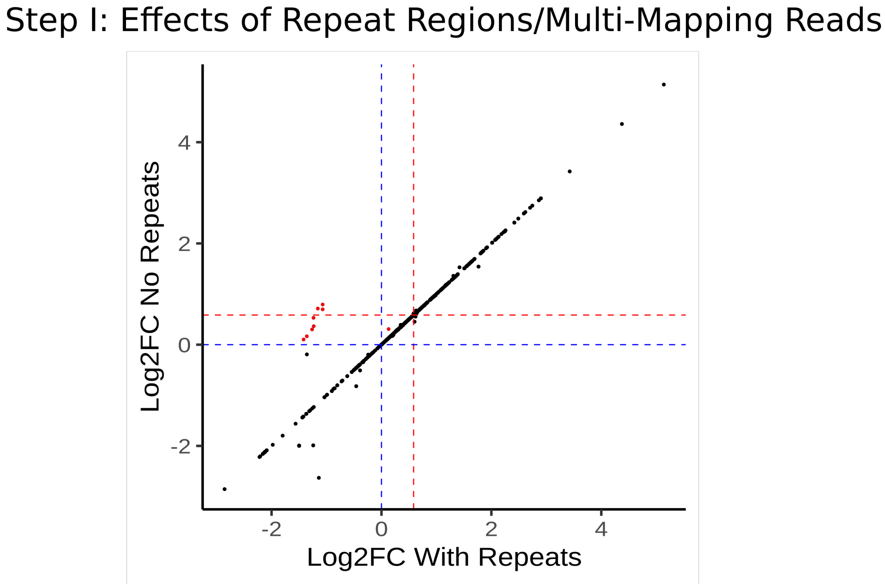


Supplementary Figure 5: **Effects of genomic repeats and multi-mapping reads on fold change estimates (T21 vs D21, RNA-seq).** Read counts were generated for all genes on chromosome 21, from bam files including genomic repeats and multi-mapping reads (x-axis), and bam files excluding these reads (y-axis). Nine genes show a reduced fold change estimation when repeat reads are included (highlighted in red). Red dashed lines indicate 1.5x fold change. Blue dashed lines indicate 1.0 fold change. The rRNA genes are highly transcribed from several locations within the genome (rDNA copies are located on chr13, chr14, chr15, chr21, chr22). Therefore, the fold change calculated when including multi-mapped reads is likely incorrect.

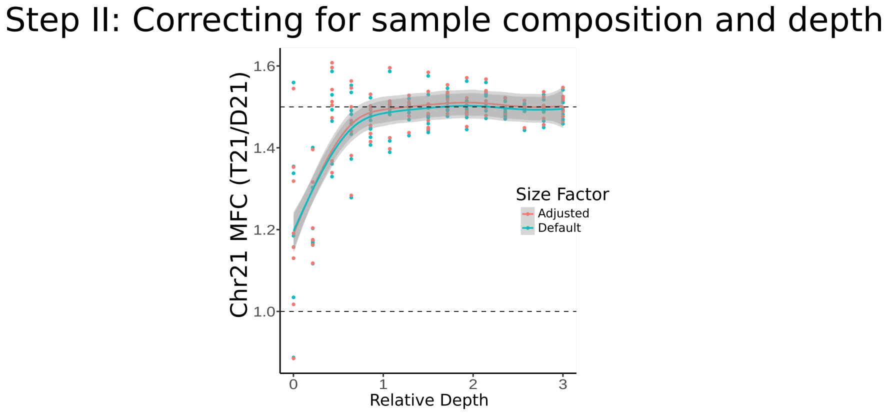


Supplementary Figure 6: **Fold change estimations in simulated data sets using two different size factor calculations.** Default (teal line): chromosome 21 genes are included in size factor calculation. Adjusted (red line): chromosome 21 genes are excluded from size factor calculation. In general, the differences in fold change estimates between these two approaches is minimal.

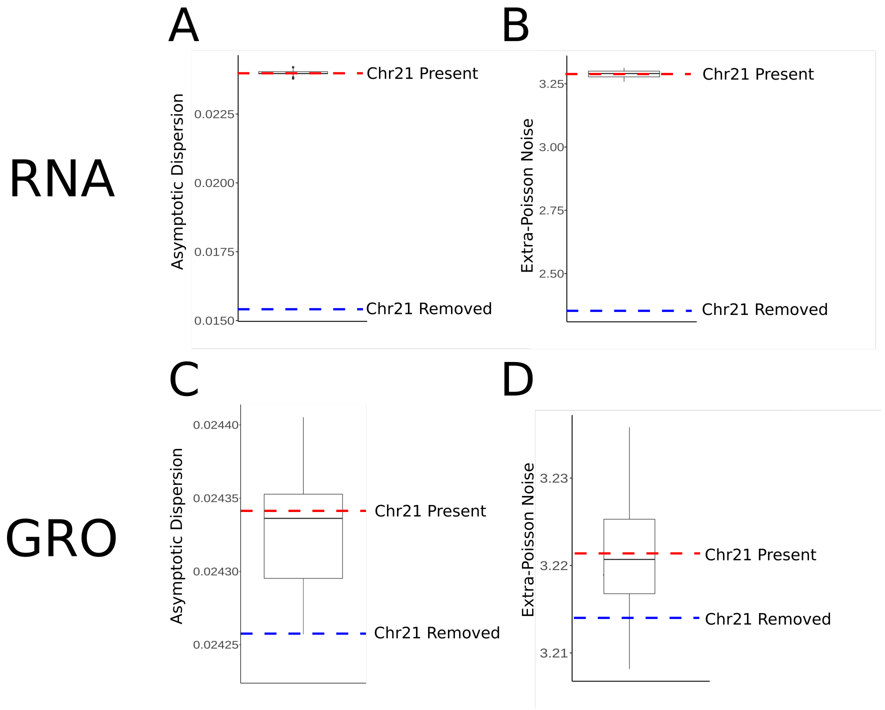


Supplementary Figure 7: **Dispersion fitting estimates calculated by DESeq2 with and without chromosome 21 genes.** Estimates are calculated including chromosome 21 genes (red) and excluding chromosome 21 genes (blue) for both RNA-seq (top) and GRO-seq (bottom). Specifically: (A) Asymptotic dispersion estimates in RNA-seq data, (B) extra-Poisson noise estimates in RNA-seq, (C) asymptotic dispersion estimates in GRO-seq data, and (D) extra-Poisson noise estimates in GRO-seq data. Additionally, an equivalent number of random non-chromosome 21 genes were excluded as a control (boxplots), showing that both values are slightly higher when chr21 is included. See Materials and Methods for information about how these values are estimated.

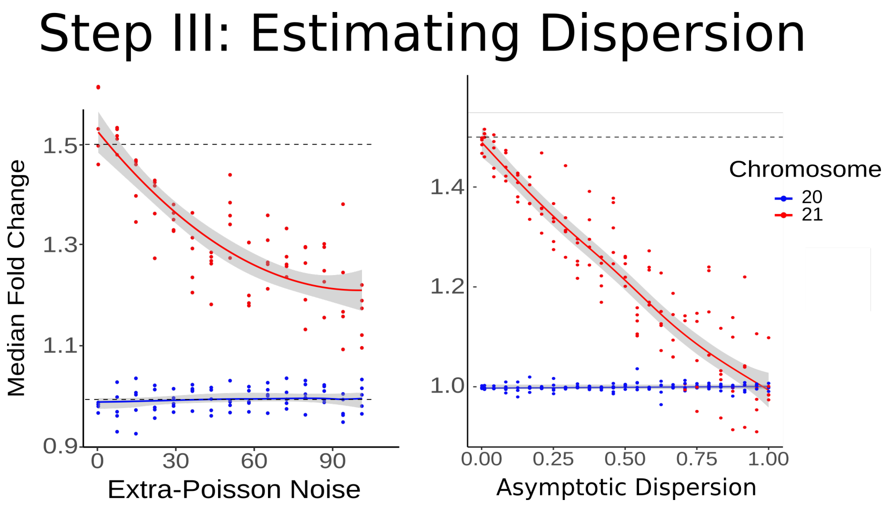


Supplementary Figure 8: **Scatterplots indicating the effects of increasing dispersion parameters on median fold change calculation in simulated data.** (Left) Effects of increasing extra-Poisson noise (asymptotic dispersion = 0.01). (Right) Effects of increasing asymptotic dispersion (extra-Poisson noise = 1). See Materials and Methods for details on simulating data.

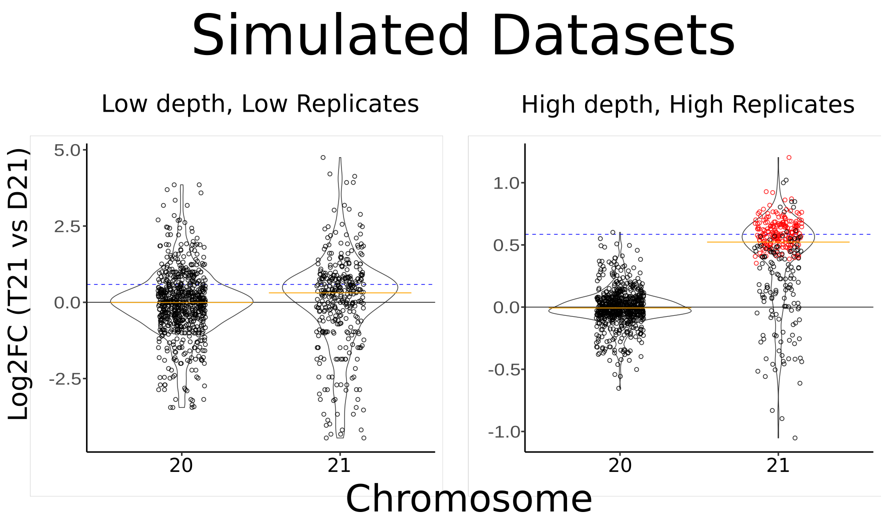


Supplementary Figure 9: **Fold-change distributions of simulated T21 and D21 data sets with high dispersion, varying the depth and replication number of the samples.** Simulations used high dispersion estimates (asymptotic dispersion=.08, extra-Poisson noise=8) and depth was changed relative to our D21 RNA-seq data (see also Supplemental Fig 1). Left: low depth (1x) and low replication (n=3). Right: high depth (3x) and high replication (n=12).

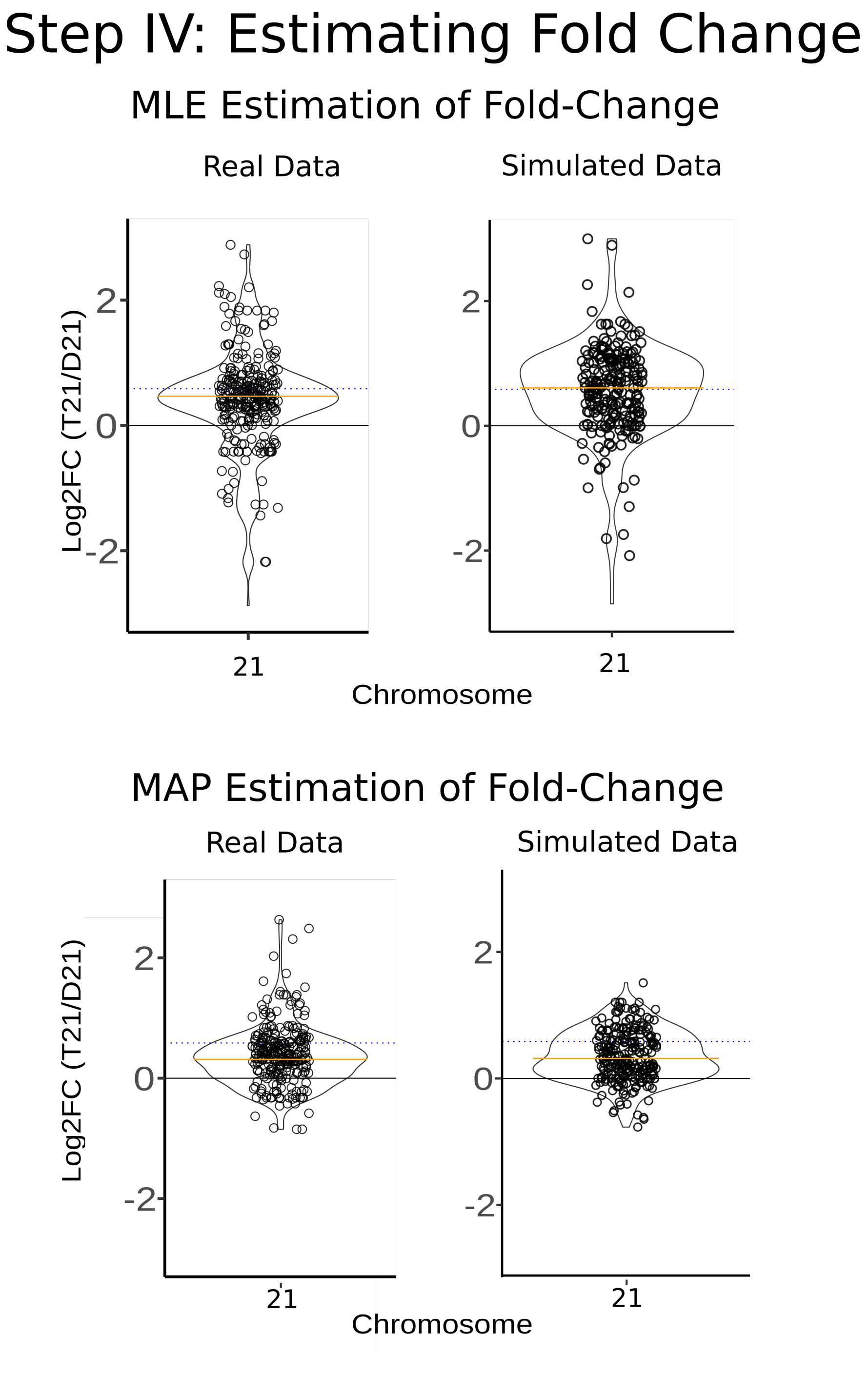


Supplementary Figure 10: **Violin plots of the fold change of chromosome 21 genes in real RNA-seq and simulated data, using maximum a posteriori estimates of fold change.** Due to fold change shrinkage, median fold change estimates are further pushed towards 0 with MAP estimation (MLE: RNA-seq MFC: 1.41, Simulated data MFC: 1.52; MAP: RNA-seq MFC: 1.24, Simulated data MFC: 1.21).

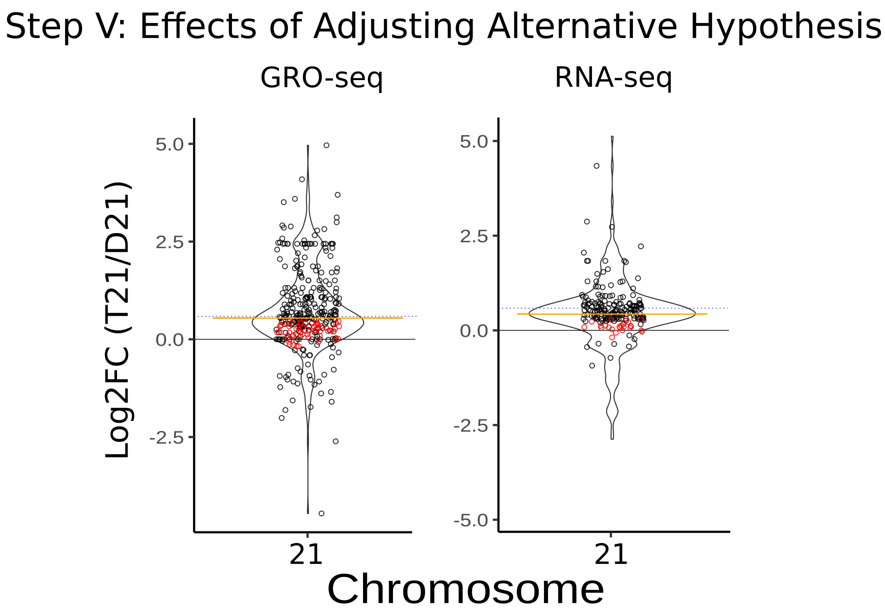


Supplementary Figure 11: **Differential analysis with adjusted alternative hypothesis (|LFC| < log2(1.5)).** Significant genes are those which are significantly below the expected value of 1.5 (dotted line). Left: GRO-seq, 56 significant genes. Right: RNA-seq, 20 significant genes

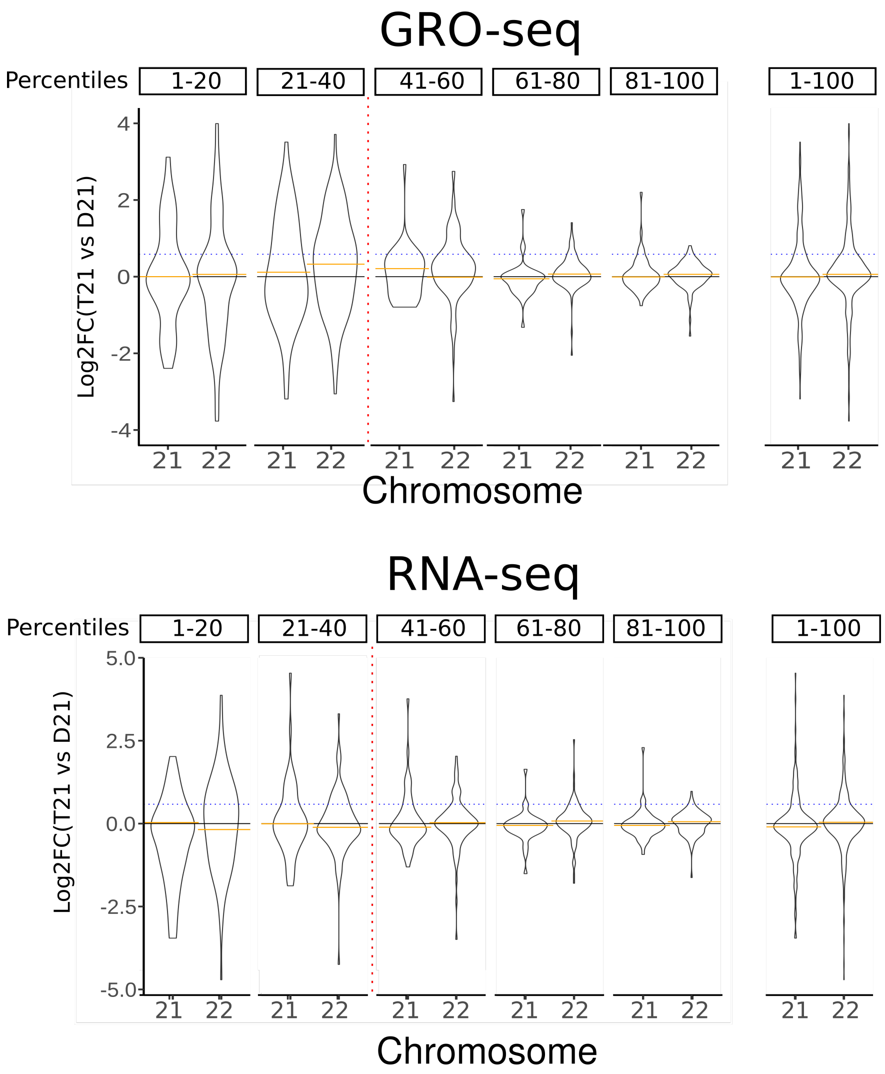


Supplementary Figure 12: **Fold change estimations (T21 vs D21) across expression levels in GRO-seq (top) and RNA-seq (bottom), after using a trisomy-aware pipeline.** Genes which fell below the red dotted line (percentiles to the left of the dotted line) are considered lowly expressed and subject to high variance, and are thus excluded from downstream analysis. See also Supplemental Fig 4 for fold change estimations without a trisomy-aware pipeline.

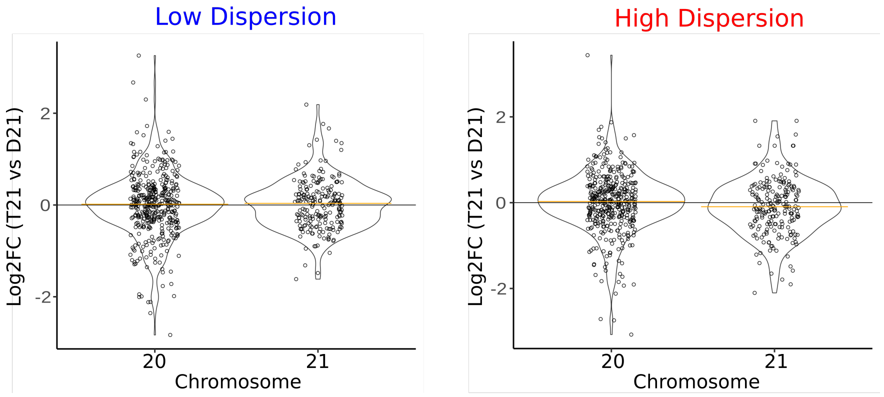


Supplementary Figure 13: **Fold-change estimates of simulated T21 and D21 data sets, using low and high dispersion estimates.** Fold-change estimates were adjusted to account for trisomy, as in Fig 2G. Low dispersion: a=.01, b=1. High dispersion: a=.05, b=30.

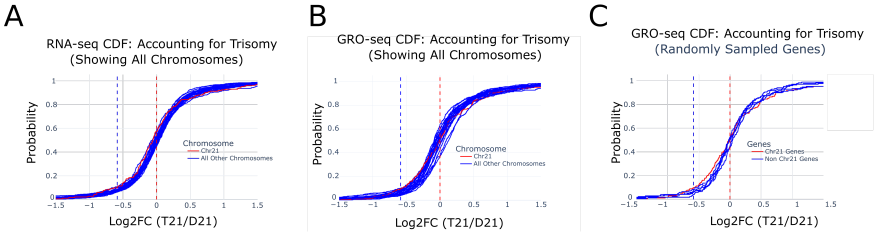


Supplementary Figure 14: **Cumulative distribution plots of fold changes in trisomy aware RNA-seq and GRO-seq analyses.** Vertical dotted red line indicates a log2 fold change of 0. Vertical dotted blue line indicates a log2 Fold change of -log2(1.5). (A) CDF of RNA-seq fold changes on all chromosomes (blue solid lines) with chromosome 21 indicated in red. (B) Same as (A), but using GRO-seq data. (C) CDF of GRO-seq fold changes using chromosome 21 genes (red) and a random subsample of an equivalent number of non-chromosome 21 genes (blue) which normalizes for size of chromosome.

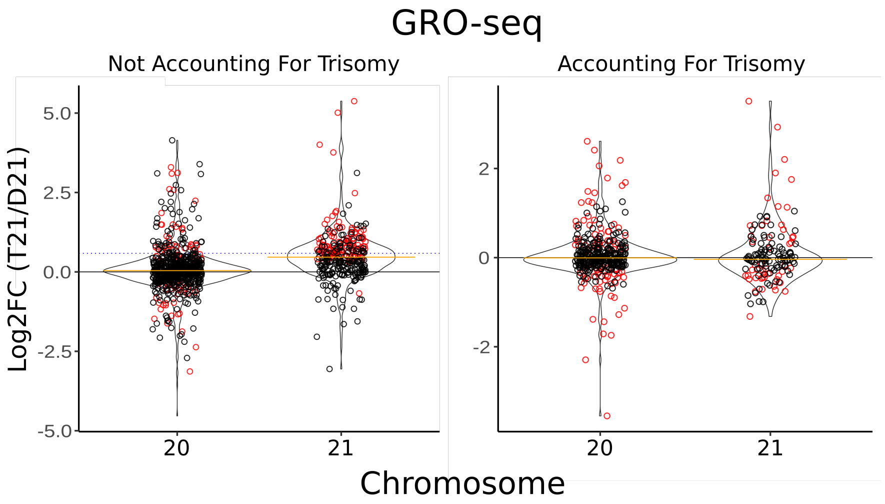


Supplementary Figure 15: **Violin plots depicting fold change estimates for chromosome 20 and 21 genes between T21 and D21 GRO-seq datasets.** (Left) Default analysis not accounting for the ploidy of the samples. (Right) Adjusted analysis correcting for ploidy differences between the samples.

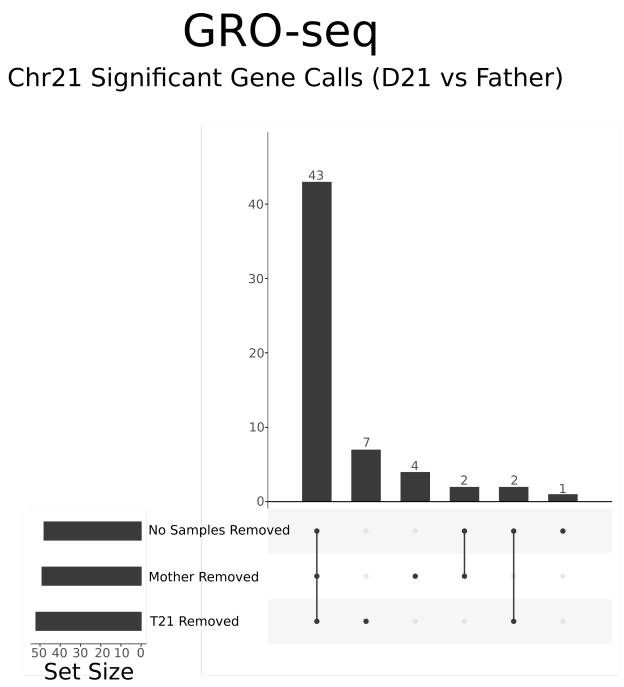


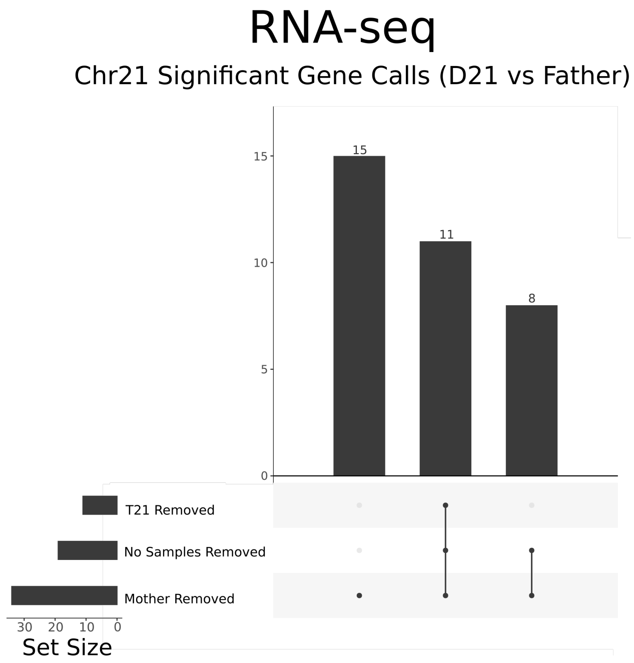


Supplementary Figure 16: **UpSet plots indicating overlap of significant gene calls on chromosome 21 comparing two disomic individuals.** We compare the D21 brother to the father and identify chromosome 21 encoded statistically significant genes (padj<.01). Three cases shown: we normalize to the entire family (no samples removed), remove the T21 sample or the mother sample is removed (as a control).

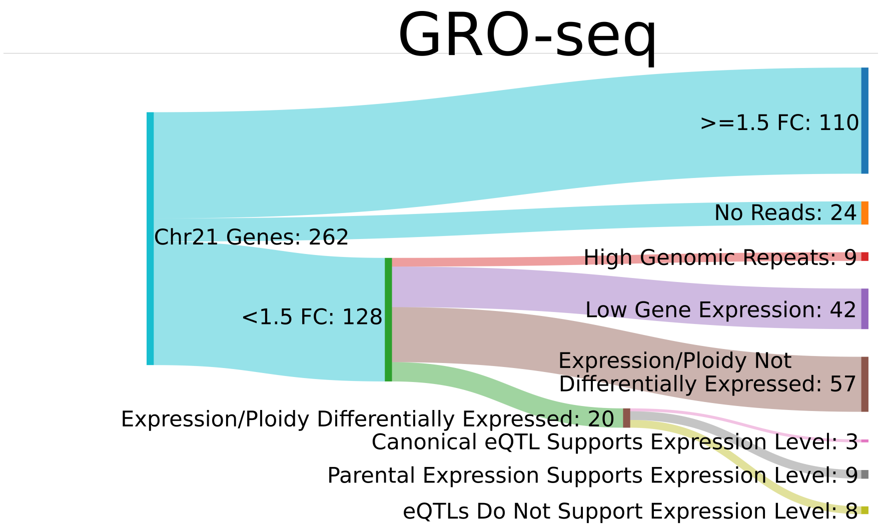


Supplementary Figure 17: **Sankey diagram indicating explanations of fold changes for chromosome 21 genes in GRO-seq (for RNA-seq, see Fig. 3D**. Nearly all genes with fold change lower than 1.5 can be explained by technical factors, leaving only 8 genes whose expression is below expectation that are not accounted for by genetic variation.
